# Tissue-specific responses to TFAM and mtDNA copy number manipulation in prematurely ageing mice

**DOI:** 10.1101/2024.11.14.623694

**Authors:** Laura S Kremer, Guanbin Gao, Giovanni Rigoni, Roberta Filograna, Mara Mennuni, Rolf Wibom, Ákos Végvári, Camilla Koolmeister, Nils-Göran Larsson

## Abstract

Somatic mitochondrial DNA (mtDNA) mutations are heavily implicated as important drivers of ageing and age-related diseases. Their pathological effect can be partially counteracted by increasing the absolute amount of wild-type mtDNA via moderately upregulating TFAM, a protein important for mtDNA packaging and expression. However, strong TFAM overexpression can also have detrimental effects as it results in hypercompaction of the mtDNA and subsequent impairment of mtDNA gene expression. In this study, we have experimentally addressed the propensity of moderate TFAM modulation to improve the premature ageing phenotypes of mtDNA mutator mice, carrying random mtDNA mutations. Surprisingly, we detect tissue-specific endogenous compensatory mechanisms acting in mtDNA mutator mice which largely affects the outcome of TFAM modulation. Accordingly, moderate overexpression of TFAM can have both negative and beneficial effects in different tissues of mtDNA mutator mice. We see a similar behavior for moderate TFAM reduction, which improves brown adipocyte tissue homeostasis, while other tissues are largely unaffected. Our findings highlight that regulation of copy number and gene expression of mtDNA is complex and cause tissue-specific effects that should be considered when modulating TFAM levels. Additionally, we suggest that TFAM is not the sole determinant of mtDNA copy number in situations where oxidative phosphorylation (OXPHOS) is compromised but other important players must be involved.

## Introduction

Ageing in animals is widely believed to be caused by accumulation of multiple types of damage and, from an evolutionary point of view, there has been little selective pressure to maintain efficient repair and maintenance of tissues at a post-reproductive, old age^1^. Although the decline of multiple pathways plays an important role^2^, the decrease of mitochondrial function is a prominent feature of ageing^3,4^. In particular, somatic mutagenesis of mtDNA is heavily associated with mammalian ageing^4^. Notably, low levels of evenly distributed mtDNA mutations may not influence the ageing process^5,6^, but rather the spatially restricted accumulation of pathogenic mtDNA mutations to high levels by clonal expansion in individual cells^7^. In ageing humans, such accumulation of mtDNA mutations result in focal respiratory chain dysfunction in a subset of cells in various tissues, e.g., in the brain, heart, skeletal muscle, and colonic crypts^8–11^. While it was long unclear whether mtDNA mutations are a mere consequence of the ageing process or contribute to cell loss and organ dysfunction, the causal role of somatic mtDNA mutations in ageing was established using genetic mouse models. In the mtDNA mutator mouse model, the expression of an exonuclease-deficient catalytic subunit of mitochondrial DNA polymerase (POLG^D257A^; genotype: *Polg*^mut/mut^) leads to extensive accumulation of mtDNA mutations causing a variety of premature ageing symptoms that drastically shorten life span^12,13^. However, it should be noted that the importance of mtDNA as a driver of ageing varies in metazoans and seems to not affect the life span in short-lived organisms. In *Caenorhabditis elegans,* life span is not affected if mtDNA replication is abrogated by knockout of *Polg*^14^, and in *Drosophila melanogaster,* a massive accumulation of somatic mutations of mtDNA caused by the expression of an exonuclease-deficient version of POLG does not limit life span^15^. The somatic mtDNA mutations associated with mammalian ageing are mostly generated during the extensive mtDNA replication that occurs during embryogenesis and postnatal development and thereafter undergo somatic segregation during adult life^16^. It is possible that the limited number of cells and cell divisions in some metazoans, like *D. melanogaster*, are insufficient to create the clonal expansion of mtDNA mutations and mosaic respiratory chain deficiency in the adult organism to limit life span. In contrast, somatic mtDNA mutations and mosaic respiratory chain deficiency seem to be ubiquitous findings in old mammals, including humans, and are experimentally linked to the ageing process through studies in the mtDNA mutator mouse strains^4,12,13^. Similar to ageing humans, most somatic mtDNA mutations in the mtDNA mutator mouse are formed during embryogenesis^17^ and clonal expansion of mutated mtDNA has been reported in some organs, e.g., heart^12^ and colonic crypts^18^. Importantly, the mtDNA mutator mouse model not only clarified the direct link between mtDNA mutations and ageing phenotypes, but it is also a valuable tool to develop strategies to mitigate the ageing process. For example, increasing mitochondrial mass by overexpressing the peroxisome proliferator-activated receptor γ coactivator-1α (PGC-1α), an important regulator of mitochondrial biogenesis, ameliorated the skeletal muscle and heart phenotype of the mtDNA mutator mouse^19^. However, PGC-1α is also involved in numerous other cellular processes and aberrant activation can be deleterious, emphasizing the need for alternative strategies to treat the natural ageing process^20–22^. A more direct approach to improving the phenotypes of the mtDNA mutator mouse is to increase the absolute amount of mtDNA molecules by upregulation of the total mtDNA copy number. While this does not affect the fraction of mutated mtDNA, the increase in the absolute number of wildtype mtDNA segments raises the likelihood that sufficient functional gene products can be made to enable sufficient OXPHOS function^23,24^. Modulation of mtDNA copy number can be achieved by genetically altering the levels of mitochondrial transcription factor A (TFAM), which packages mtDNA into nucleoids. Heterozygous knockout of *Tfam* in wild-type mice results in ∼50% decrease of mtDNA levels, whereas moderate overexpression of *Tfam* leads to ∼50% increase in mtDNA levels^25,26^. Importantly, moderate decrease or increase in TFAM levels mainly affects mtDNA copy number but does only mildly or not at all alter mtDNA gene expression, respiratory chain function or mitochondrial mass in mice without mtDNA mutations. Employing this strategy to increase mtDNA levels in the mtDNA mutator mouse revealed a rescuing effect on the early-onset male infertility phenotype at four months of age^24^. Similarly, a beneficial impact was also evident in the m.C5024T tRNA^Ala^ mouse model where moderate overexpression of TFAM and the associated mtDNA increase improved the cardiomyopathy phenotype in aged mice^27^. Despite these largely positive outcomes of upregulating mtDNA copy number via moderate TFAM overexpression, care must be taken as strong overexpression of TFAM is known to lead to respiratory chain deficiency in skeletal muscle and shorten the life span in mice^26^. It is now clear that the TFAM-to-mtDNA ratio determines the compaction of the nucleoid, and *in vivo* and *in vitro* experiments have established that a high degree of nucleoid compaction will shut off mtDNA expression^26,28,29^.

Here, we investigated whether mtDNA copy number modulation via moderate TFAM alteration impacts the ageing phenotypes in mtDNA mutator mice. Surprisingly, a moderate increase of TFAM levels did not translate to a proportional modulation of the mtDNA copy number, as previously observed in mice with wild-type mtDNA or in the tRNA^Ala^ mouse model. Instead, the mtDNA mutator mouse shows a trend towards endogenous upregulation of mtDNA copy number in some tissues and experimental modulation of TFAM will be added to this increase to result in tissue-specific effects with varying TFAM-to-mtDNA ratios that impact mtDNA gene expression in different ways. In colon, TFAM overexpression reduced mtDNA gene expression, negatively affecting OXPHOS and leading to a marked increase in expression of *Methylenetetrahydrofolate dehydrogenase 2* (*Mthfd2)*, a marker for mitochondrial dysfunction. In contrast, in brown adipose tissue (BAT), a decrease in TFAM levels normalized *Uncoupling protein 1* (*Ucp1)* expression. In summary, mtDNA copy number regulation is more complex than suggested by previous studies^23–27^ and the TFAM-to-mtDNA ratio seems to be finely tuned in a tissue-specific manner. Alterations of the ratio can have different outcomes in different tissues and attempts to increase TFAM levels to counteract pathologies caused by mitochondrial dysfunction must therefore be carefully evaluated.

## Results

### Moderate alterations of TFAM levels only mildly affect pathology in aged mtDNA mutator mice

To experimentally test the impact of mtDNA copy number modulation on the premature ageing phenotypes of mtDNA mutator mice, we generated hemizygous Polg^-/mut^ mice with moderately increased (*Tfam*^+/OE^) or decreased (*Tfam*^+/-^) TFAM levels. We first created heterozygous Polg knockout (*Polg*^+/-^) females that also carry either an *Tfam*^+/OE^ or *Tfam*^+/-^ allele, and subsequently mated those females to heterozygous mtDNA mutator (*Polg*^+/mut^) males (Supp. Fig. 1). Employing this mating scheme ensures that the resulting mice (*Polg*^-/mut^) carry a substantial load of *de novo* generated somatic mtDNA mutations and prevent that they inherit maternally transmitted mtDNA mutations from their mothers, which otherwise would aggravate the phenotype^30^. Therefore, in the studied *Polg*^-/mut^ mice, all mtDNA mutations are generated *de novo* during embryonic and postnatal life. Both the “classical” mtDNA mutator mice (*Polg*^mut/mut^)^12,13^ and the mtDNA mutator mice generated in this study (*Polg*^-/mut^) develop profound premature ageing phenotypes at the age of 35 weeks. The mtDNA mutator mice (*Polg*^-/mut^) had a markedly reduced body weight at the age of 35 weeks and increasing TFAM levels (*Polg*^-/mut^; *Tfam*^+/OE^) led to a statistically significant further, albeit slight, reduction of body weight (Fig. 1A). In contrast, reduction of TFAM expression (*Polg^-/mut^; Tfam^+/-^*) did not have any statistically significant effect on the reduced body weight of mtDNA mutator mice (Fig. 1A). The organ-to-body weight ratios were largely unaffected by variations in TFAM expression, with the exception of testis. In testis, we observed a clear rescue effect in mtDNA mutator mice with increased TFAM levels (*Polg*^-/mut^; *Tfam*^+/OE^) resulting in a testis to body weight ratio similar to wildtype mice, whereas reduced TFAM levels (*Polg*^-/mut^; *Tfam*^+/-^) exacerbated the phenotype (Fig. 1, B-D), consistent with our previous study^24^. These results demonstrate that TFAM modulation does not grossly impact most ageing phenotypes of the mtDNA mutator mouse.

**Figure 1:**
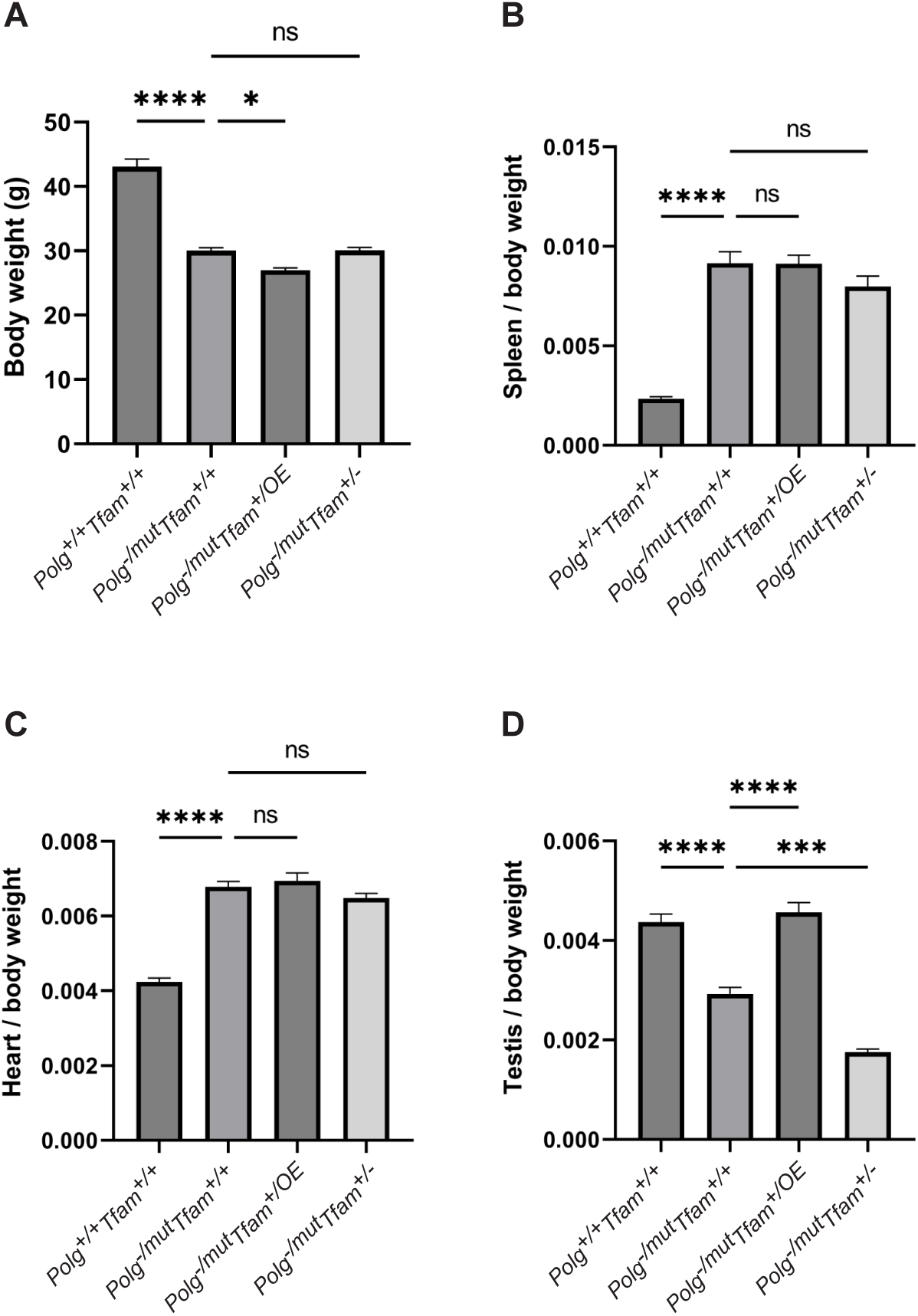
TFAM modulation does not rescue the decreased body weight or the increased spleen/body weight and heart/body weight ratio. A) Body weight in g. B) Spleen to body weight ratio (g/g). C) Heart to body weight ratio (g/g). D) Testis to body weight ratio (g/g). n≥7. Data are represented as mean ± SEM; *p< 0.05; **p< 0.01; ***p< 0.001; ns: non-significant.

### Tissue-specific effects on mtDNA copy number and TFAM-to-mtDNA ratios

The beneficial effect of moderate TFAM overexpression on the testis phenotype, and the absence of any rescue effect on the increased heart and spleen phenotype of *Polg*^-/mut^; *Tfam*^+/OE^ mice prompted us to perform further characterization of mtDNA expression in different tissues. To this end, we analyzed the relative mtDNA copy number by qPCR using three different probes, i.e., *NADH-ubiquinone oxidoreductase chain 1* (*mt-Nd1*), *ATP synthase subunit a* (*mt-Atp6*) and *Cytochrome b* (*mt-CytB*). It should be noted that the mtDNA mutator mice carry linear deleted mtDNA molecules spanning the major arc of mtDNA^12^ (Fig. 2A). These linear deletions are continuously formed replication intermediates and represent around 30% of the total mtDNA in all investigated tissues^12,31^. *Mt-Nd1* hybridizes to a genomic region corresponding to the minor arc of the mtDNA and is hence a good proxy for full-length mtDNA, whereas *mt-Atp6* and *mt-CytB* hybridize to genomic regions in the major arc of mtDNA and therefore are good indicators of total mtDNA levels, including deleted mtDNA molecules (Fig. 2A). The mtDNA mutator mice spontaneously upregulated the total mtDNA copy number in several tissues (Fig. 2 B-F, Supp. Fig. 2A), similar to what has previously been shown for mice harboring a point mutation in the tRNA^Ala^ gene^27^. Surprisingly, increased TFAM levels did not cause any additional relative mtDNA copy number change in either liver or heart of *Polg*^-/mut^; *Tfam*^+/OE^ mice and only a very subtle difference in the colon (Fig. 2B-D). The qPCR results for the liver were independently confirmed by Southern blot analyses (Supp. Fig. 2, B and C). In contrast, increased TFAM levels caused an additional relative mtDNA copy number increase in brown adipose tissue and spleen of *Polg*^-/mut^; *Tfam*^+/OE^ mice compared to mtDNA mutator mice (Fig. 2 E,F).

**Figure 2:**
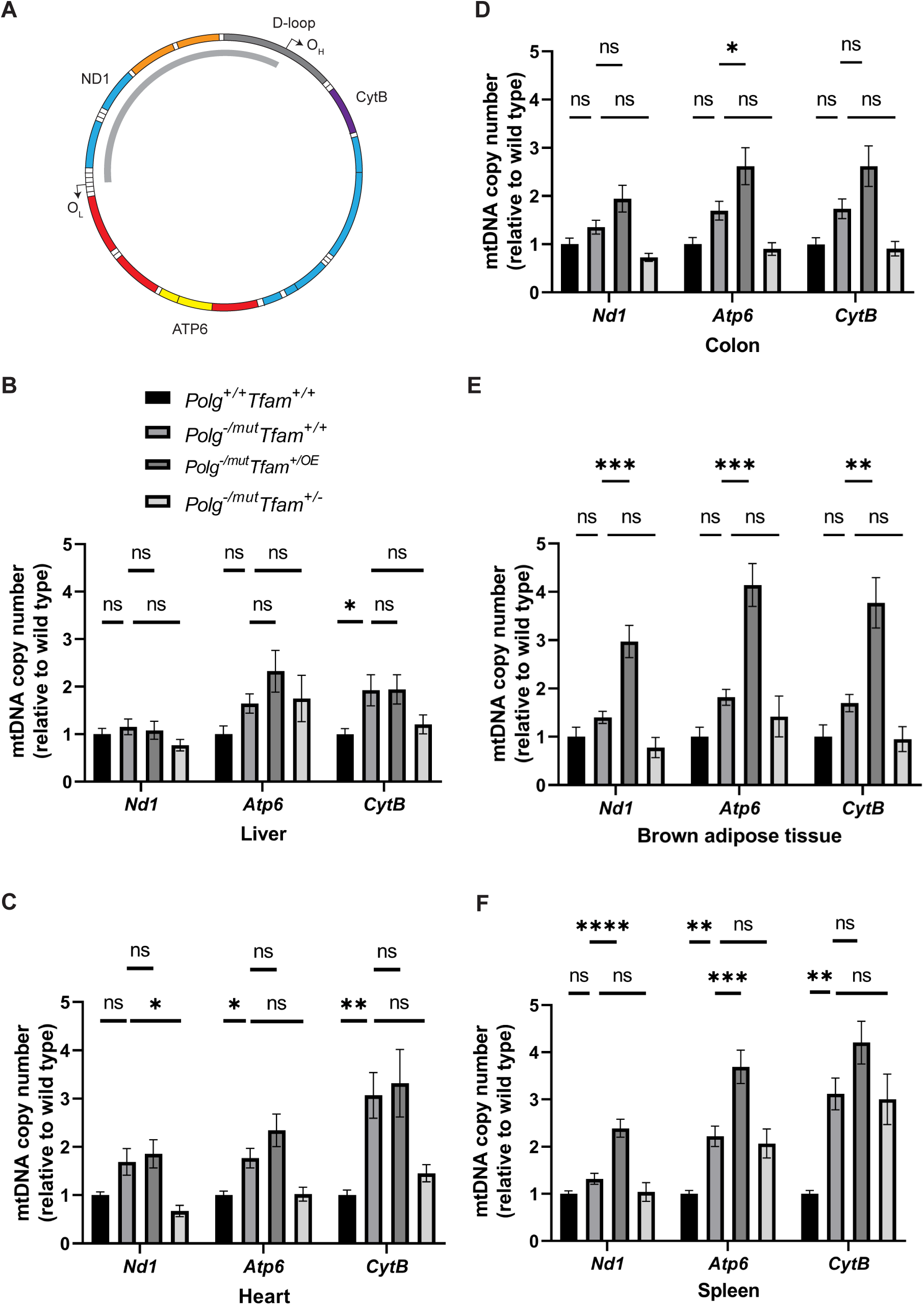
Modulation of TFAM expression affects mtDNA copy number in a tissue-specific manner. A) Schematic of the mtDNA highlighting the position of the probes used for mtDNA copy number analysis by qPCR. The deleted mtDNA region is indicated with a grey arc. B-F) Relative mtDNA copy number quantification (*Nd1*/*18S*, *Atp6*/*18S*, *Cytb*/*18S*) in B) liver. C) heart D) colon E) brown adipose tissue and F) spleen. n≥5. Data are represented as mean ± SEM; *p< 0.05; **p< 0.01; ***p< 0.001; ns: non-significant.

Changes in TFAM levels without a concomitant change in mtDNA levels are potentially problematic, because the TFAM-to-mtDNA ratio will be impacted to affect nucleoid compaction, which, in turn, impacts mtDNA expression^26,28,29^. We therefore proceeded to calculate the relative TFAM-to-mtDNA ratios in liver, heart, colon, spleen, and brown adipose tissue. Compared to wild-type mice, *Polg*^-/mut^; *Tfam*^+/+^ mice had a moderately elevated TFAM-to-mtDNA ratio in the heart and a drastically decreased ratio in the spleen. TFAM overexpression in the mtDNA mutator mice (*Polg*^-/mut^; *Tfam*^+/OE^ mice) resulted in a marked additional increase of the TFAM-to-mtDNA ratio in the liver and an additional increase in the heart in comparison with *Polg*^-/mut^; *Tfam*^+/+^ mice, whereas no major effect was observed in the other tissues (Table 1). When the TFAM levels instead were decreased (*Polg*^-/mut^; *Tfam*^+/-^ mice), the TFAM-to-mtDNA ratios were decreased in liver, colon, and BAT in comparison with *Polg*^-/mut^; *Tfam*^+/+^ mice, whereas no additional change was seen in heart (Table 1).

**Table 1.**
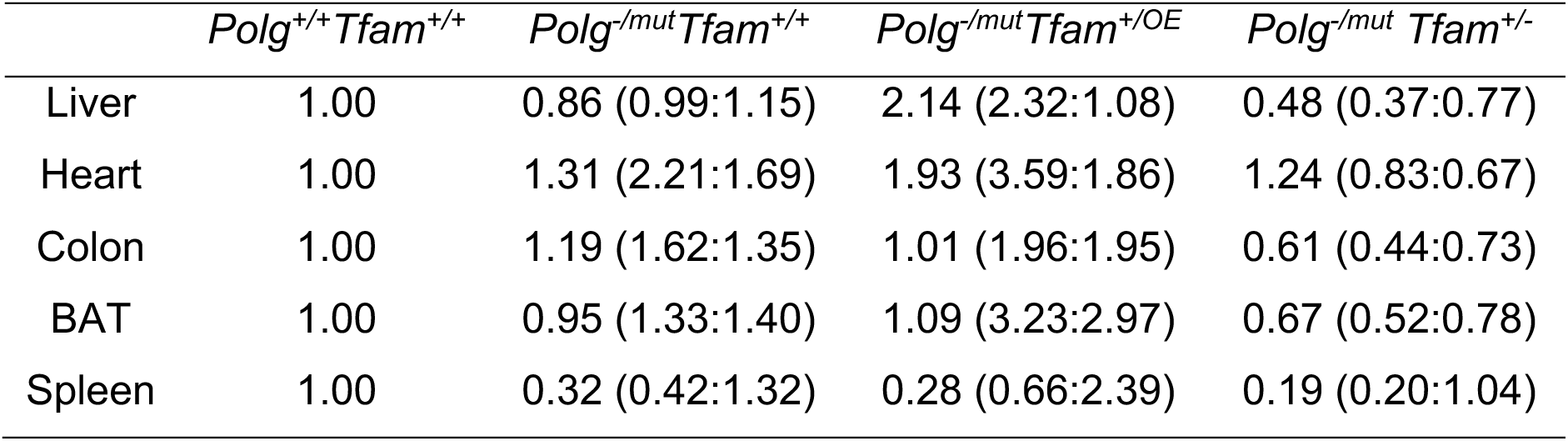
Relative TFAM-to-mtDNA ratios in different tissues. The TFAM-to-mtDNA ratio was calculated from normalized TFAM protein levels (n=2) and normalized mtDNA levels (n=5) as determined by the ND1 probe. The respective values for the normalized TFAM and mtDNA levels are indicated in parentheses. BAT: brown adipose tissue.

Taken together, these findings show that modulation of TFAM expression in the severely affected mtDNA mutator mouse has tissue-specific effects on TFAM-to-mtDNA ratios, which is well known to influence nucleoid compaction^26^. We therefore proceeded to investigate the pathophysiological effects of TFAM modulation in different organs of the mtDNA mutator mouse. We hypothesized that in tissues where TFAM overexpression increases the TFAM-to-mtDNA ratio, the mtDNA expression will be reduced and negatively impact tissue function. In contrast, we speculated that in tissues where elevated TFAM levels increase mtDNA copy number, the mitochondrial function would improve, in line with previous results^24,27^.

### Moderate TFAM overexpression impairs mtDNA expression in the liver of mtDNA mutator mice

In the liver, TFAM overexpression resulted in a decrease of *mt-Atp6* and *mt-Cytb* transcript levels (Fig. 3A) as predicted by the increased TFAM-to-mtDNA ratio. The decrease in gene expression was also evident on the protein level as the steady-state levels of several subunits of the OXPHOS complexes were further decreased as compared to *Polg*^-/mut^; *Tfam*^+/+^ mice (Fig. 3B). Of note, we did not only observe a reduction in levels of proteins encoded by the mtDNA, e.g., Cytochrome c oxidase subunit 1 (COX1) and Cytochrome c oxidase subunit 2 (COX2) of complex IV, but also of OXPHOS subunits encoded by the nuclear DNA, e.g., NADH:Ubiquinone Oxidoreductase Subunit B8 (NDUFB8) of complex I. This is an expected secondary effect as mtDNA gene expression is necessary for the stability of nucleus-encoded OXPHOS complex subunits^32^. The levels of the ATP Synthase F1 Subunit Alpha (ATP5A) protein of complex V (ATP synthase) were largely unaffected, which goes well in line with previous reports showing that impaired mtDNA expression leads to the formation of a stable subcomplex containing the F1 subunit of complex V of the OXPHOS system^32^. We detected increased levels of the complex II subunit Succinate Dehydrogenase Complex Iron Sulfur Subunit B (SDHB). Complex II is exclusively nuclear encoded and a compensatory increase upon impaired mitochondrial gene expresson has been observed before^32^.

**Figure 3:**
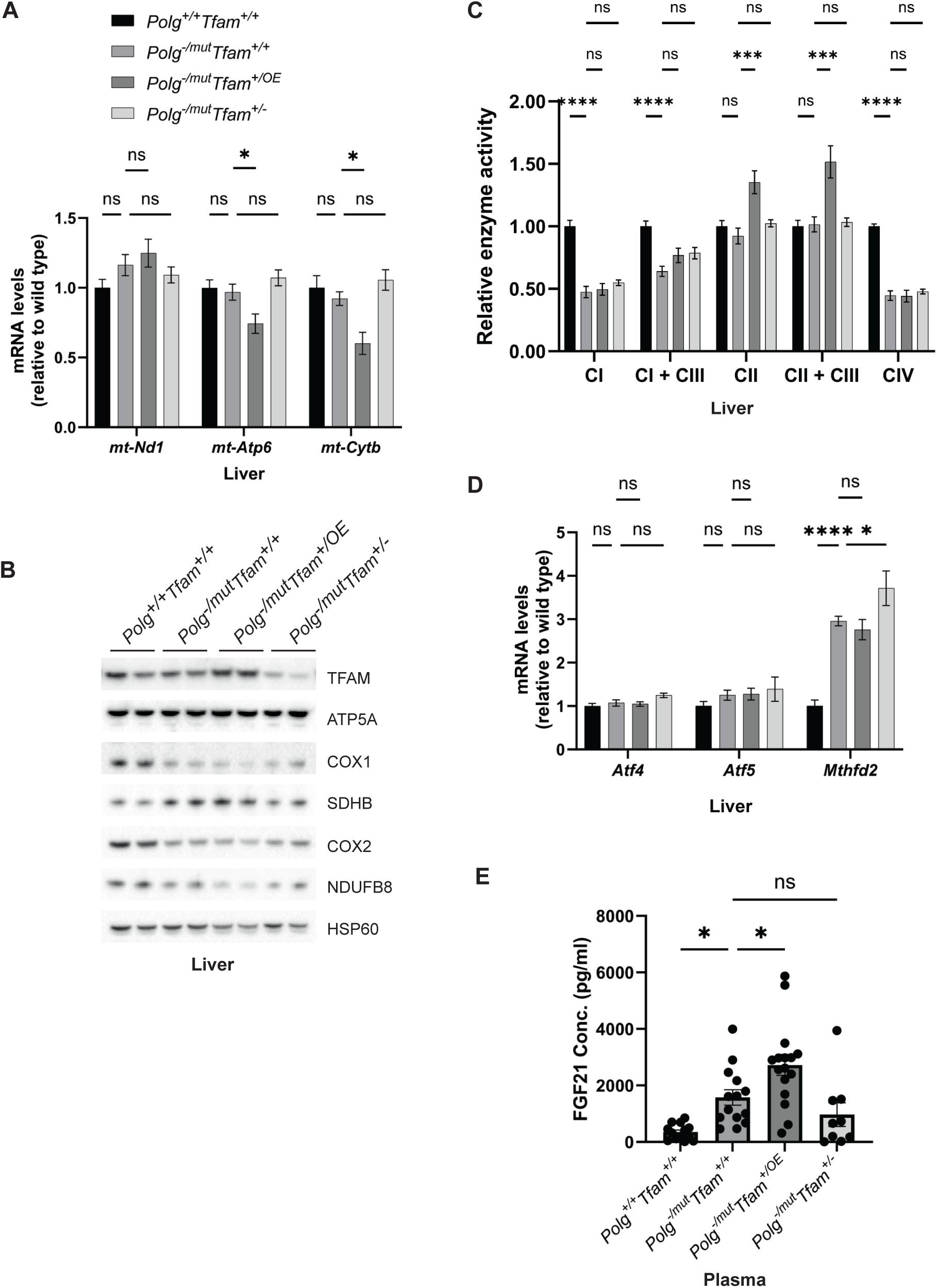
Moderate TFAM overexpression negatively impacts mtDNA gene expression and tissue physiology in liver and correlates with increased FGF2 levels. A) Relative expression levels of mtDNA encoded transcripts (*Nd1*/μ*-actin*, *Atp6*/μ*-actin*, *Cytb*/μ*-actin*) measured by RT-qPCR in liver. n≥7. Data are represented as mean ± SEM; *p< 0.05; **p< 0.01; ***p< 0.001; ns: non-significant. B) Western blot analysis of steady-state levels of mitochondrial proteins in liver. C) Relative enzyme activities of OXPHOS complexes measured by spectrophotometry in liver mitochondria. n ≥ 6 Data are represented as means ± SEM; *P < 0.05; **P < 0.01; **p< 0.01; ***P < 0.001; ns: non-significant. D) Relative expression levels of mitochondrial stress markers (*Atf4*/μ*-actin*, *Atf5*/μ*-actin*, *Mthfd2*/μ*-actin*) measured by RT-qPCR in liver. n ≥7. Data are represented as means ± SEM; *P < 0.05; **p< 0.01; ***P < 0.001, ns: non-significant E) Quantification of FGF21 levels in plasma measured by ELISA. n *≥ 9.*Data are represented as means ± SEM; *P < 0.05; **p< 0.01; ***P < 0.001; ns: non-significant.

We proceeded to measure the enzyme activities of individual OXPHOS complexes in liver mitochondria (Fig. 3C). The complex I and complex IV activities were reduced to about 50% in *Polg*^-/mut^; *Tfam*^+/+^ mice in comparison with wild-type mice (Fig. 3C). However, we did not see any further alteration of the reduced enzyme activities induced by TFAM overexpression or reduced TFAM expression (Fig. 3C). Interestingly, we detected a significant increase in complex II and complex II + complex III activity upon TFAM overexpression, which can partially be explained by the increased complex II protein levels we oberseved in *Polg*^-/mut^; *Tfam*^+/OE^ mice (Fig. 3, B and C). The *Polg*^-/mut^; *Tfam*^+/+^ mice had normal expression of *Activating Transcription Factor 4* (*Atf4)* and *Activating Transcription Factor 5 (Atf5)* in liver, whereas the expression of *Mthfd2* was markedly increased (Fig. 3D). We have previously reported that increased *Mthfd2* expression is a sensitive marker of mitochondrial dysfunction and that *Mthfd2* expression increases as at OXPHOS dysfunction progresses^32^. The finding of increased *Mthfd2* expression in *Polg*^-/mut^; *Tfam*^+/+^ liver is thus consistent with the observed OXPHOS dysfunction (Fig. 3, C and D). This was unaltered by TFAM overexpression.

We also observed an increase of Fibroblast Growth Factor 21 (FGF21) levels in plasma of *Polg^-/mut^; Tfam^+/+^* mice pointing to a negative effect on energy homeostasis (Fig. 3E). Interestingly, the plasma levels of FGF21 were further increased in *Polg*^-/mut^; *Tfam^+/OE^*mice (Fig. 3E). These findings are consistent with the proposed action of FGF21 as a starvation hormone and may reflect the progression of body weight loss in *Polg^-/mut^; Tfam^+/+^* and additional body weight reduction in *Polg*^-/mut^; *Tfam^+/OE^* mice (Fig. 1A).

In summary, moderate TFAM overexpression does not improve the liver phenotype of mtDNA mutator mice. On the contrary, elevating TFAM had a negative effect on mtDNA gene expression (Fig. 3A and 3B), which could be linked to the increased TFAM-to-mtDNA ratio (Table 1) causing tighter compaction of the mitochondrial nucleoid. The mtDNA mutator mouse has abundant point mutations in mtDNA that will affect tRNAs and rRNAs thus impairing mitochondrial translation, as well as abundant point mutations causing amino acid substitutions in the protein coding genes of mtDNA thus causing dysfunction or impaired stability of respiratory chain complexes. The impaired mtDNA gene expression caused by increased nucleoid compaction poses an additional burden on the already compromised OXPHOS dysfunction in mtDNA mutator mice. The marked additional increase in FGF21 levels in *Polg*^-/mut^; *Tfam^+/OE^*mice indicates that TFAM overexpression has a negative impact on liver physiology in mtDNA mutator mice.

### Modulation of TFAM does not impact the cardiomyopathy phenotype of mtDNA mutator mice

In the heart of *Polg*^-/mut^; *Tfam*^+/+^ mice, we detected strongly reduced levels of mtDNA-encoded transcripts (Fig. 4A). This can potentially be explained by the substantial increase of TFAM protein levels in these animals (Fig. 4B), which, given the absence of a corresponding increase in full-length mtDNA, results in a higher TFAM-to-mtDNA ratio and thus probably causes reduced mtDNA expression (Table 1). Importantly, TFAM modulation did not have any effect on the reduced transcript levels or the diminished protein levels of OXPHOS complexes in mtDNA mutator mice (Fig. 4A, B).

**Figure 4:**
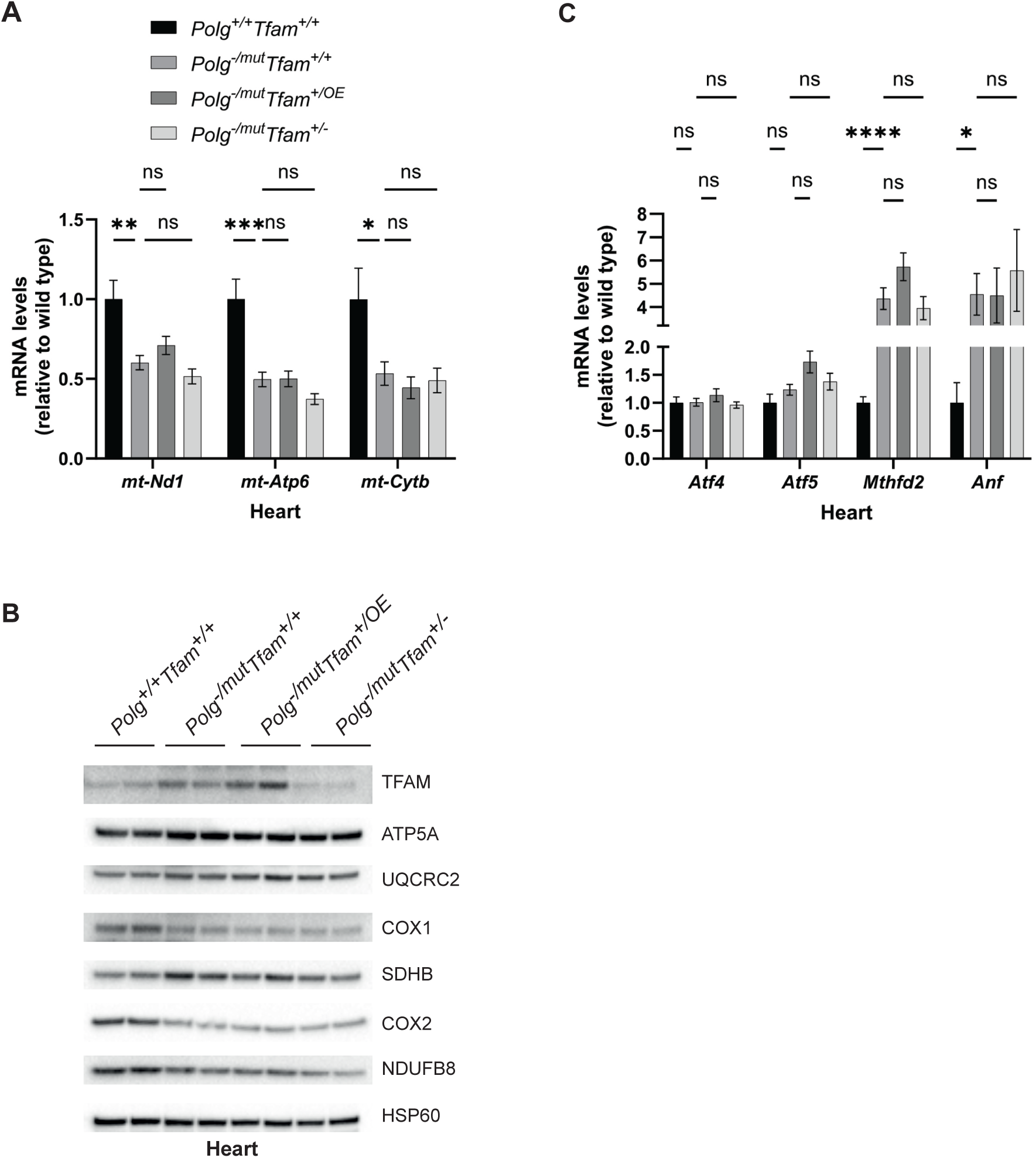
Alteration of TFAM expression does not affect the heart phenotype of mtDNA mutator mice. A) Relative expression levels of mtDNA encoded transcripts (*Nd1*/μ*-actin*, *Atp6*/μ*-actin*, *Cytb*/μ*-actin*) measured by RT-qPCR in heart. n *≥ 7*. Data are represented as mean ± SEM; *p< 0.05; **p< 0.01; ***p< 0.001; ns: non-significant. B) Western blot analysis of steady-state levels of mitochondrial proteins in heart. C) Relative expression levels of mitochondrial stress markers (*Atf4*/μ*-actin*, *Atf5*/μ*-actin*, *Mthfd2*/μ*-actin*, *Anf*/μ*-actin*) measured by RT-qPCR in heart. n *≥ 7*. Data are represented as mean ± SEM; *p< 0.05; **p< 0.01; ***p< 0.001; ns: non-significant.

Consistent with the increased heart-to-body-weight ratio of *Polg*^-/mut^; *Tfam*^+/+^ mice (Fig. 1C), the expression of *Natriuretic Peptide A* (*Anf*), a marker for heart failure, and *Mthfd2*, as described above a marker for OXPHOS dysfunction, were increased, whereas there was no change in mRNA levels for *Atf4* and *Atf5*. (Fig. 4 C). The levels of *Atf4*, *Atf5*, *Mthfd2* and *Anf* mRNAs were not impacted by reduced or increased expression of TFAM (Fig. 4C). These findings are in agreement with the observation that TFAM overexpression does not rescue the increased heart-to-body weight ratio observed in mtDNA mutator mice (Fig. 1C).

All in all, these results demonstrate that the mtDNA mutator hearts upregulate TFAM protein levels, likely as a compensatory mechanism. However, the increased TFAM levels do not lead to a concomitant increase in mtDNA levels, but instead result in an increased TFAM-to-mtDNA ratio and reduced steady-state levels of mitochondrial transcripts. TFAM overexpression does not further affect this endogenous compensatory response and causes no additional increase in mtDNA levels. Thus, neither moderate TFAM overexpression nor reduced TFAM expression affect the heart phenotype of mtDNA mutator mice.

### Increased TFAM levels do not rescue OXPHOS dysfunction in colon of mtDNA mutator mice

In colon, we found an increase of levels of mtDNA-encoded mRNAs in *Polg*^-/mut^; *Tfam*^+/+^ mice (Fig. 5A). This response was largely ablated upon moderate TFAM overexpression. In line with this, levels of key OXPHOS subunits were further reduced in *Polg*^-/mut^; *Tfam^+/^*^OE^ mice in comparison to *Polg*^-/mut^; *Tfam*^+/+^ mice (Fig. 5B). However, this did not lead to a further deterioration of levels of respiratory chain complexes on blue-native PAGE gels (Fig. 5C) or respiratory chain enzyme activities (Fig. 5D), possibly due to their already extremely low levels in *Polg*^-/mut^; *Tfam*^+/+^ mice. We found a strong induction of the *Atf5* and *Mthfd2* mRNAs in colon of *Polg*^-/mut^; *Tfam*^+/+^ mice. Upon TFAM overexpression, *Mthfd2* mRNA expression levels were further increased (Fig. 5E).

**Figure 5:**
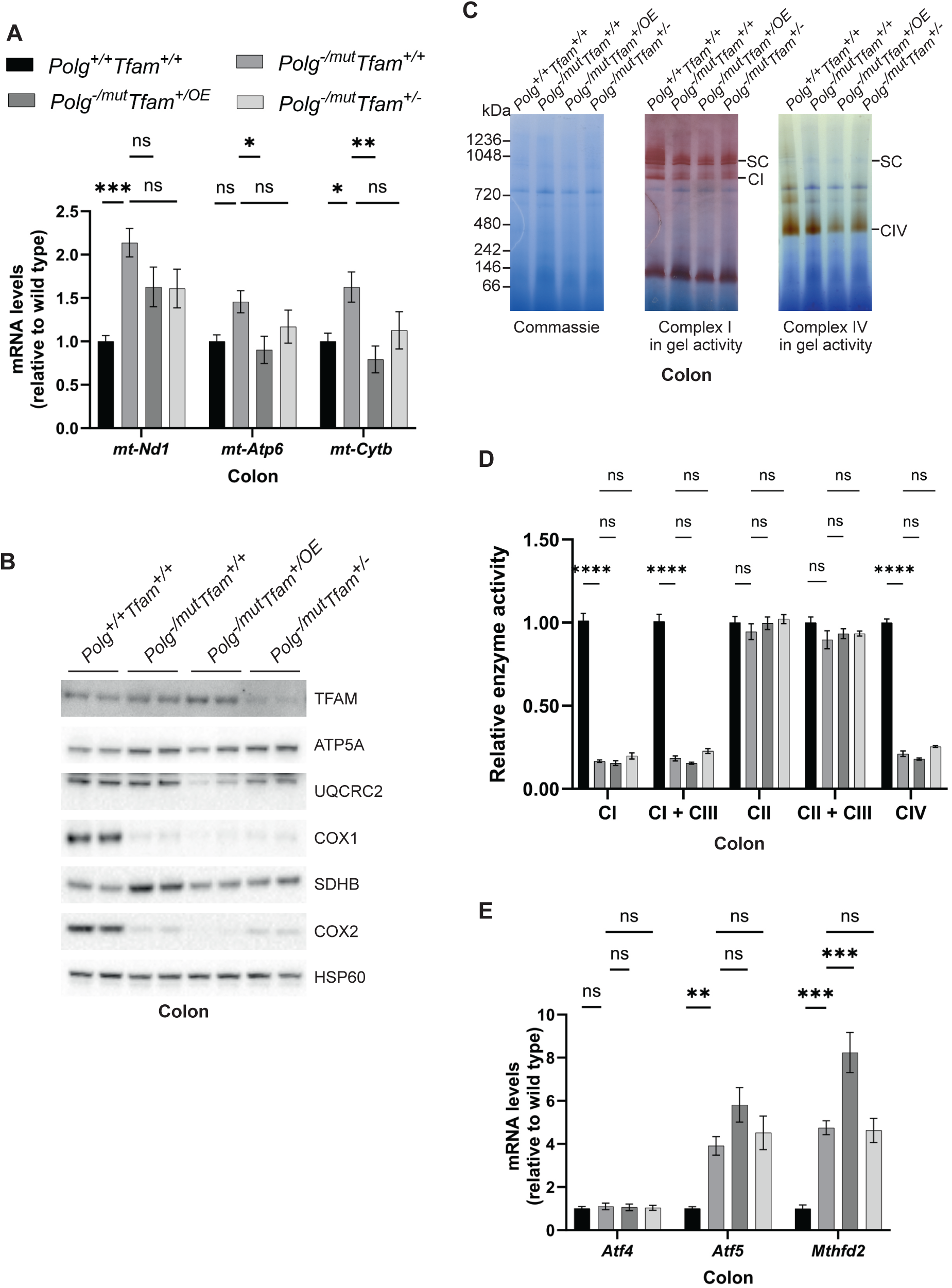
Increased TFAM levels do not rescue the reduced OXPHOS function in colon of mtDNA mutator mice. A) Relative expression levels of mtDNA encoded transcripts (*Nd1*/μ*-actin*, *Atp6*/μ*-actin*, *Cytb*/μ*-actin*) measured by RT-qPCR in colon. n ≥7. Data are represented as mean ± SEM; *p< 0.05; **p< 0.01; ***p< 0.001; ns: non-significant. B) Western blot analysis of steady-state levels of mitochondrial proteins in colon. C) BN-PAGE and in-gel activities of complex I and complex IV activities in mitochondrial protein extracts from mouse colon. Coomassie staining of the gel is shown to indicate equal loading. SC, Supercomplexes. D) Relative enzyme activities of OXPHOS complexes measured by spectrophotometry in colon mitochondria. n ≥3. Data are represented as mean ± SEM; *p< 0.05; **p< 0.01; ***p< 0.001; ns: non-significant. E) Relative expression levels of mitochondrial stress markers (*Atf4*/μ*-actin*, *Atf5*/μ*-actin*, *Mthfd2*/μ*-actin*) measured by RT-qPCR in colon. n ≥9. Data are represented as mean ± SEM; *p< 0.05; **p< 0.01; ***p< 0.001; ns: non-significant.

In summary, the responses in the colon have similarities to the responses in the liver as TFAM overexpression results in a reduction of mtDNA-encoded transcripts in mtDNA mutator mice. This reduction likely adds to the already drastic decrease of protein levels of several subunits of the OXPHOS complexes. The strongly induced expression of *Mthfd2* suggests that instead of having a beneficial effect, TFAM overexpression results in a deterioration of colon physiology.

### TFAM downregulation rescues Ucp1 expression in brown adipose tissue of mtDNA mutator mice

TFAM overexpression in BAT resulted in a significant increase in mtDNA copy number and did hence not alter the TFAM-to-mtDNA ratio (Fig. 2E and Table 1). In line with this finding, the levels of mtDNA-encoded transcripts (Fig. 6A) and OXPHOS subunits (Fig. 6B) did not change. In contrast, the reduced TFAM expression in *Polg*^-/mut^; *Tfam*^+/-^ mice led to increased steady-state levels of mtDNA-encoded transcripts (Fig. 6A), which correlate well with the decrease in TFAM-to-mtDNA ratios in BAT of these mice. The levels of OXPHOS subunits in *Polg*^-/mut^; *Tfam*^+/-^ mice were only mildly increased or not changed (Fig. 6B). UCP1 protein levels are reduced in mtDNA mutator mice and TFAM overexpression did not influence the UCP1 levels in BAT. Surprisingly, the reduced TFAM expression in *Polg*^-/mut^; *Tfam*^+/-^ mice resulted in elevated UCP1 protein levels (Fig. 6B). Consistent with this result, the expression of *Ucp1* and *Cell Death Inducing DFFA Like Effector A* (*Cidea)* mRNAs, which are markers for the thermogenic competence of BAT, were strongly induced in *Polg*^-/mut^; *Tfam*^+/-^ mice compared to mtDNA mutator mice (Fig. 6C). These findings argue that restoration of mtDNA expression may be important for maintaining the differentiated state of BAT in *Polg*^-/mut^ mice or that homeostatic mechanisms induced by altered function of other tissues affect nuclear gene expression in BAT.

**Figure 6:**
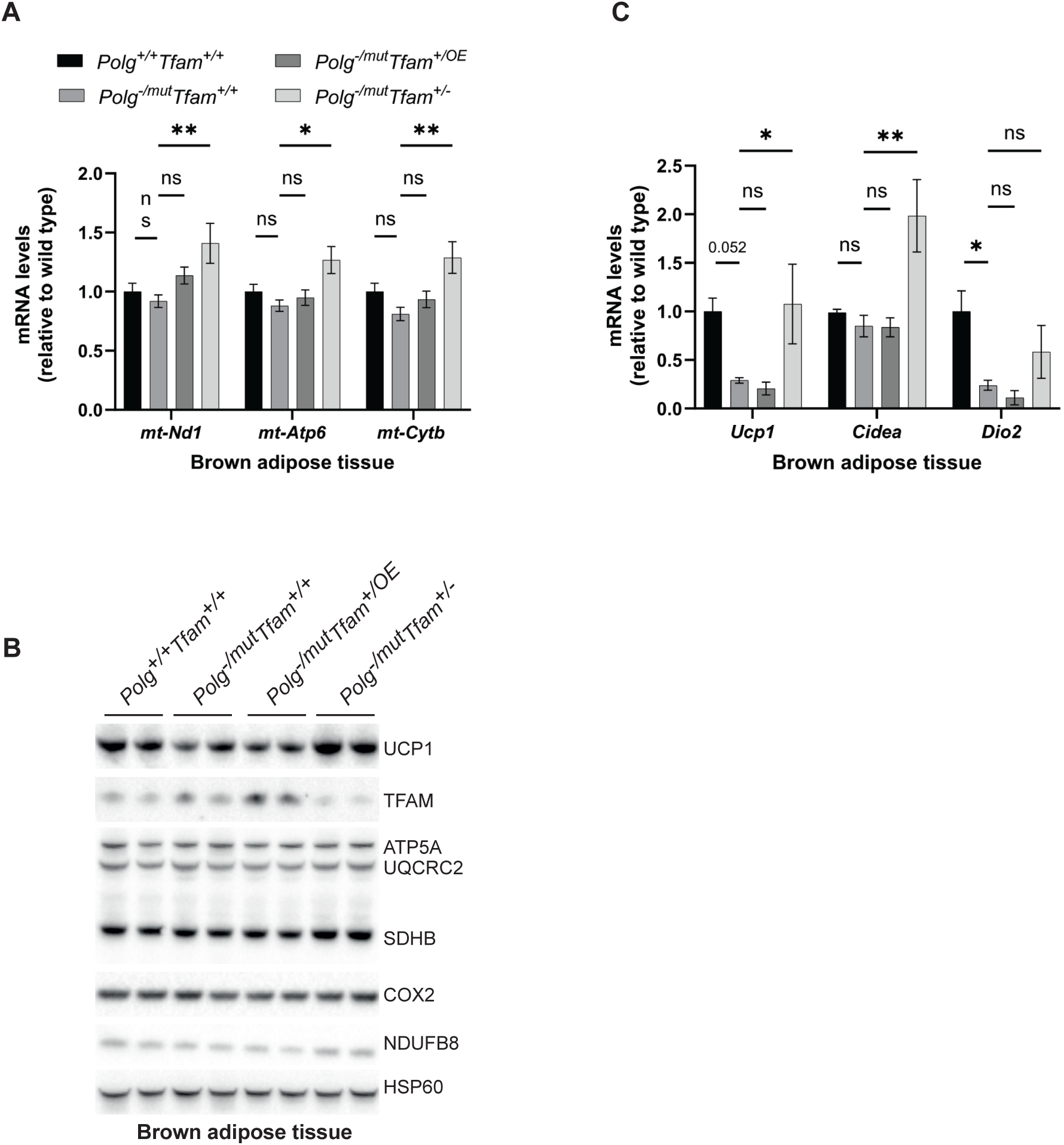
Reduction of TFAM levels in brown adipose tissue has beneficial effects. A) Relative expression levels of mtDNA encoded transcripts (*Nd1*/μ*-actin*, *Atp6*/μ*-actin*, *Cytb*/μ*-actin*) measured by RT-qPCR in brown adipose tissue (BAT). N ≥5. Data are represented as mean ± SEM; *p< 0.05; **p< 0.01; ***p< 0.001; ns: non-significant. B) Western blot analysis of steady-state levels of UCP1 and mitochondrial proteins in BAT. C) Relative expression levels of brown adipose stress markers (*Ucp1*/μ*-actin*, *Cidea*/μ*-actin*, *Dio2*/μ*-actin*) measured by RT-qPCR in BAT. n≥5. Data are represented as mean ± SEM; *p< 0.05; **p< 0.01; ***p< 0.001; ns: non-significant.

Taken together, our data show that moderate TFAM overexpression in BAT causes an increase in mtDNA copy number without affecting mtDNA expression. While this could potentially lead to a beneficial effect on BAT, we did not observe any rescue effect when assessing transcript and protein levels of key factors important for BAT function. In contrast, reduced TFAM levels led to a considerable burst in mtDNA gene expression which surprisingly ameliorated the expression of markers for the thermogenic competence of BAT, hence likely positively affecting BAT function.

### Upregulation of TFAM does not impact mitochondrial gene expression in mtDNA mutator spleen but impacts cytokine levels

In spleen of *Polg*^-/mut^; *Tfam*^+/+^ mice, levels of mtDNA-encoded transcripts were substantially increased (Fig. 7A). This is likely connected to the decrease in TFAM protein levels in these animals (Fig. 7B) which is not accompanied by a corresponding increase in mtDNA levels but instead result in a markedly decreased TFAM-to-mtDNA ratio (Table 1). Despite the elevated transcript levels in spleen of mtDNA mutator mice, the levels of the complex IV subunit COX2 were drastically decreased (Fig. 7B). TFAM overexpression, which further increased mtDNA copy number in spleen (Fig. 2F), did not affect the TFAM-to-mtDNA ratio (Table 1) or mtDNA gene expression (Fig. 7A and B). This resembles the previously reported situation in testis and heart where moderate TFAM overexpression in young *Polg*^mut/mut^ mice or aged tRNA^Ala^-mutant mice, respectively, exerted a beneficial effect by increasing the absolute amount of mtDNA without affecting the mtDNA mutation load^24,27^. To assess whether the increased mtDNA copy number in *Polg*^-/mut^; *Tfam*^+/OE^ mice led to amelioration of the premature ageing phenotype in the spleen, we measured the levels of several cytokines in plasma. The mtDNA mutator mice demonstrated elevated Interleukin 5 (IL-5) and C-C Motif Chemokine Ligand 2 (CCL2) cytokine levels in plasma and increased *ll5* transcript levels in spleen (Fig 7C and 7D). TFAM overexpression normalized the levels of IL-5 and CCL2 in plasma (Fig. 7C) and reduced the levels of *ll5* transcripts in spleen (Fig. 7D). The levels of a range of other cytokines in plasma did not change (Supp. Fig. 3). We determined the proportion of immune cell populations in spleen by using gene expression markers and found a significant reduction of all T helper cell lineage markers (*T-Box Transcription Factor 21* (*Tbx21), GATA Binding Protein 3 (Gata3), RAR Related Orphan Receptor C (Rorc)* and *Forkhead Box P3* (*Foxp3)*) in mtDNA mutator spleen. In contrast, a granulocyte marker (*CCAAT Enhancer Binding Protein Epsilon, Cebpe*) was increased (Fig. 7E). This finding implies a significant shift in immune cell populations in the spleen of *Polg*^-/mut^; *Tfam*^+/+^ mice. TFAM overexpression did not affect the expression of the immune cell markers in spleen. The mtDNA mutator mice have been shown to have disrupted white pulp structure in the spleen due to persistent inflammation^33^. However, the overall histology of the spleen did not reveal any apparent rescue effect by TFAM modulation (Fig. 7F).

**Figure 7:**
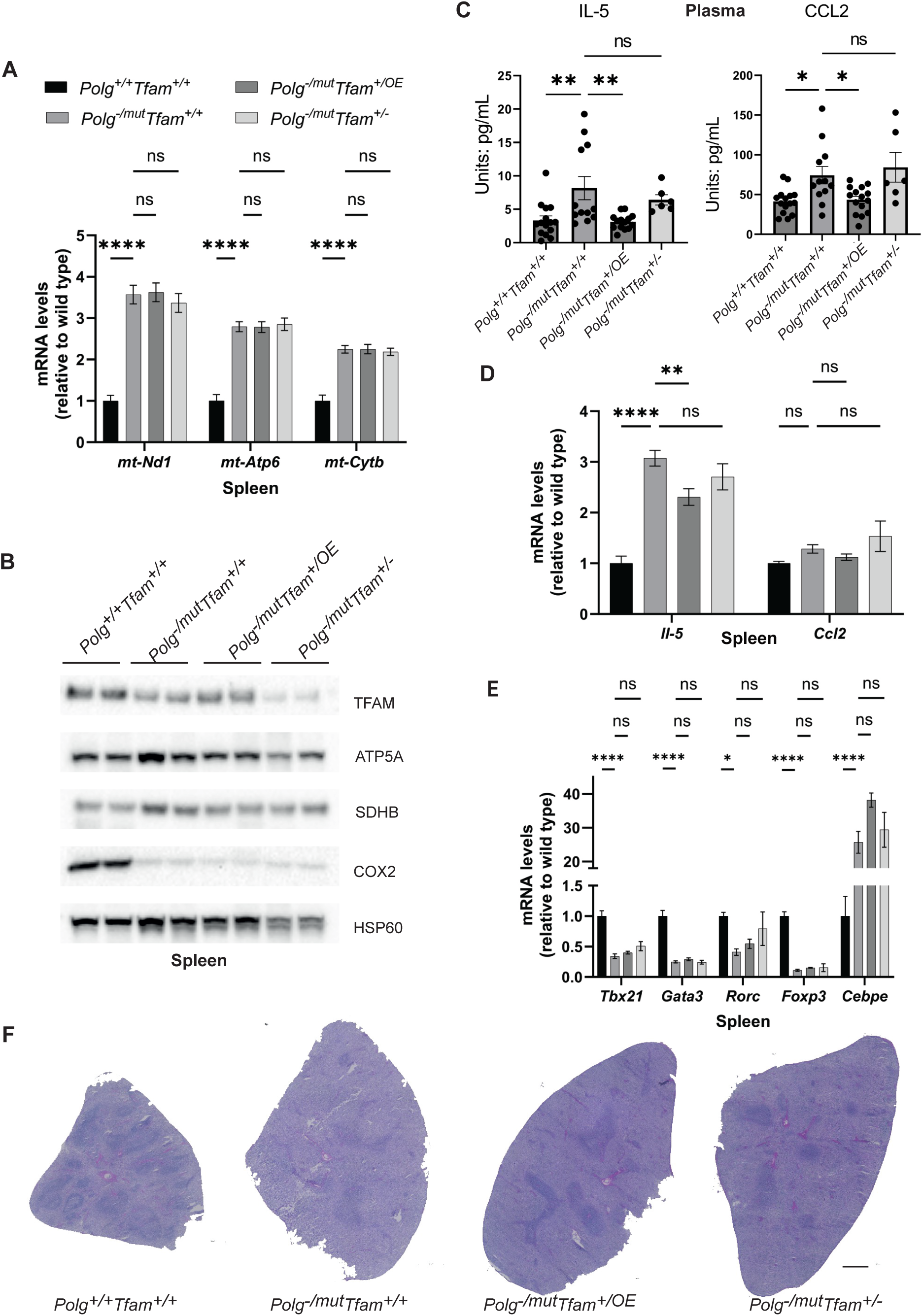
TFAM overexpression restores IL-5 and CCL2 cytokine levels in spleen of mtDNA mutator mice. A) Relative expression levels of mtDNA encoded transcripts (*Nd1*/μ*-actin*, *Atp6*/μ*-actin*, *Cytb*/μ*-actin*) measured by RT-qPCR in spleen. n ≥5. Data are represented as mean ± SEM; *p< 0.05; **p< 0.01; ***p< 0.001; ns: non-significant. B) Western blot analysis of steady-state levels of mitochondrial proteins in spleen. C) Quantification of IL-5 and CCL2 cytokine levels in plasma measured by the Mouse Cytokine/Chemokine 44-Plex Discovery Assay. n ≥6. Data are represented as mean ± SEM; *p< 0.05; **p< 0.01; ***p< 0.001; ns: non-significant. D) Quantification of *Il-5* and *Ccl2* cytokine transcript levels in spleen measured by RT-qPCR. n ≥10. Data are represented as mean ± SEM; *p< 0.05; **p< 0.01; ***p< 0.001; ns: non-significant. E) Relative expression levels of immune cell markers (*Tbx21*/μ*-actin*, *Gata3*/μ*-actin*, *Rorc*/μ*-actin*, *Foxp3*/μ*-actin*, *Cebpe*/μ*-actin*) for analyzing immune cell populations measured by RT-qPCR in spleen. n ≥5. Data are represented as mean ± SEM; *p< 0.05; **p< 0.01; ***p< 0.001; ns: non-significant. F) H&E staining of spleen sections. scale bar: 500µm.

Finally, we subjected spleen samples to tandem mass tag-based quantitative proteomic analysis to reveal any further beneficial or detrimental effects of TFAM modulation. In comparison with wild-type mice, the *Polg*^-/mut^; *Tfam*^+/+^ spleen showed upregulation of proteins involved in heme biosynthesis and reactive oxygen species (ROS) defense pathways. The former is consistent with the severe anemia present in the mtDNA mutator mice^34^, causing compensatory extramedullary haematopoiesis^35^. The latter is possibly due to altered ROS signaling which is an important player for reshuffling hematopoietic cell populations^36^. Increasing TFAM levels mitigated the induction of both of these pathways (Supp. Fig. 4). The beneficial effect can potentially be attributed to an acute improvement of OXPHOS function due to the increased levels of wildtype mtDNA segments, to a long-term proliferative advantage and hence clonal expansion of cells harboring less mutated mtDNA molecules, or a combination of both.

To summarize, TFAM elevation evokes corresponding changes of mtDNA levels in the spleen, while mtDNA gene expression is unaffected. The absolute increase of mtDNA seen in TFAM overexpressing mice leads to a positive effect on spleen physiology, in line with previous findings in testis of mtDNA mutator mice^24^.

## Discussion

We demonstrate here that even moderate changes of TFAM levels can have drastically different outcomes on the premature ageing phenotypes of mtDNA mutator mice. Depending on the tissue, increase in TFAM levels can be detrimental or beneficial. Likewise, reducing TFAM levels also shows ambivalent effects. Whereas most tissues did not seem to be affected by reduced TFAM levels, BAT gene expression was reconstituted indicating normalized physiology. This highlights the complexity and tissue-specificity of the regulation mtDNA copy number and expression and warrants careful evaluation of the therapeutic potential of TFAM alterations in different disease settings.

Somatic mutations of mtDNA are important contributors to the ageing process and are found in various common age-associated diseases including cancer, neurodegeneration, diabetes, and cardiac disorders^37^. They act by causing focal respiratory chain dysfunction, eventually compromising cell function and tissue homeostasis^8–11^. Several elegant strategies have been developed to ameliorate the accompanying pathology. One strategy has focused on increasing mitochondrial mass by overexpressing PGC-1α and has been effective in improving the skeletal muscle and heart phenotypes of the mtDNA mutator mouse^19^. However, PGC-1α levels must be carefully titrated as it is involved in several other cellular processes and as aberrant activation of PGC-1α can be harmful^20–22^. Similarly, tampering with mitochondrial mass might be problematic given the diverse functional roles of mitochondria. Another promising and more direct strategy is to increase the mtDNA copy number, and thereby the absolute amount of wild-type mtDNA molecules, by raising TFAM levels. Notably, also TFAM modulation is not without risk because mtDNA copy number does follow TFAM levels only up until a certain point, which seems to be tissue specific. Increasing TFAM levels beyond this point does not result in a further rise in mtDNA copy number, but instead increases the TFAM-to-mtDNA ratio^26^. The TFAM-to-mtDNA ratio determines nucleoid compaction, which, in turn, is crucial for mtDNA gene expression. Strong TFAM overexpression can therefore lead to impaired mtDNA expression and respiratory chain deficiency in a tissue-specific manner, and eventually shorten the life span of the mouse^26^. Nonetheless, moderate TFAM overexpression by about 50% has been shown to be well tolerated in all mouse models investigated so far^24,26,27^. Indeed, this rational also demonstrated to be effective in rescuing the early-onset male infertility phenotype in mtDNA mutator mice of 4 months of age^24^. The TFAM increase resulted in doubling of the mtDNA copy number in mouse testis. While the total mtDNA mutation load was unaltered, the increased mtDNA levels thus led to a higher absolute number of mtDNA molecules carrying segments without mutations. Likewise, increasing the absolute number of wild-type mtDNA copies via moderately increasing TFAM expression in the m.C5024T tRNA^Ala^ mouse model showed a beneficial effect on the cardiomyopathy phenotype in aged mice^27^.

We did not observe ubiquitously beneficial effects when employing moderate TFAM overexpression to alleviate the premature ageing phenotypes of mtDNA mutator mice. On the one hand, TFAM elevation in spleen led to a concomitant increase of full-length mtDNA translating into improved spleen homoeostasis. On the other hand, increasing TFAM levels in liver did not result in higher mtDNA levels but instead caused impaired mtDNA expression. The tissue-specific responses in mtDNA mutator mice differs from what was seen in mice without mtDNA mutations or in mice carrying the heteroplasmic pathogenic m.C5024T tRNA^Ala^ mutation as well as mice carrying a mtDNA deletion^23^. The response in mtDNA mutator mice resembles the tissue-specific effects seen in mice with very strong TFAM overexpression where the mtDNA levels follow TFAM levels only up to a certain point in a tissue-specific manner. The discrepancy in outcomes of moderate TFAM overexpression can potentially be explained b y the varying presence of compensatory mechanisms attributed to different levels of OXPHOS malfunction in the investigated mouse models. The m.C5024T tRNA^Ala^ mouse model harbors a heteroplasmic maternally inherited pathogenic mtDNA mutation that at high levels causes a mitochondrial translational defect which mildly affects OXPHOS protein levels and function^27,38^. In contrast, the mtDNA mutator mouse continuously generates new mtDNA mutations every time an mtDNA molecule is replicated and it is therefore much more severely affected. The abundant mtDNA mutations in the mtDNA mutator mice affects the tRNAs and rRNAs needed for mitochondrial translation and cause abundant amino acid substitutions in the mtDNA-encoded OXPHOS subunits. The OXPHOS deficiency in the mtDNA mutator mice is thus caused by a combination of reduced synthesis mtDNA-encoded OXPHOS subunits and synthesis of mtDNA-encoded OXPHOS subunits with defective function, which will reduce both the levels and the function of the OXPHOS complexes. The more pronounced pathology in mtDNA mutator mice likely explains the activation of compensatory mechanisms consistent with the intrinsic upregulation of TFAM found in the heart. As such, TFAM is necessary but not sufficient to control mtDNA levels under pathological conditions on its own, but other licensing factors seem to be limiting. While TFAM seems to be the main protein coating mtDNA, the precise stoichiometry and positioning likely alters between different cellular states, providing access for other proteins such as the mitochondrial single-stranded DNA binding protein (mtSSB)^29,39–42^. Additionally, further licensing factors could, for example, include the abundance of other proteins involved in mtDNA replication and expression and goes in line with previous studies in cultured mammalian cells as well as budding yeast^43–45^. Notably, the severity of the OXPHOS dysfunction largely differed between tissues and, accordingly, the endogenous compensatory mechanisms as well as the rescuing efficacy of TFAM modulation was tissue specific. The different vulnerability of a given tissue towards OXPHOS dysfunction caused by mtDNA mutations is likely dependent on its physiology, metabolism, proliferative character, absolute mtDNA levels, mtDNA turnover rates, the TFAM-to-mtDNA ratios and additional factors.

In summary, the mtDNA mutator mouse shows tissue-specific endogenous compensatory mechanisms in response to the continuous mutagenesis of mtDNA, including the upregulation of TFAM expression, elevated mtDNA copy number, as well as altered mtDNA gene expression. This likely limits the impact of genetic manipulation of TFAM expression and explains the variable tissue-specific consequences of altered TFAM expression on mtDNA copy number, mtDNA gene expression and tissue physiology. Importantly, this also clearly demonstrates that TFAM is not the only determinant of mtDNA levels under pathological conditions, but argues that other factors involved in mtDNA replication must play a limiting role. Identifying these additional molecular players holds the promise to more accurately intervene with the ageing process and counteract other pathological entities caused by mtDNA mutations including mitochondrial disorders, cancer, neurodegenerative diseases, diabetes, and cardiac disorders.

## Materials and Methods

### Key resources table

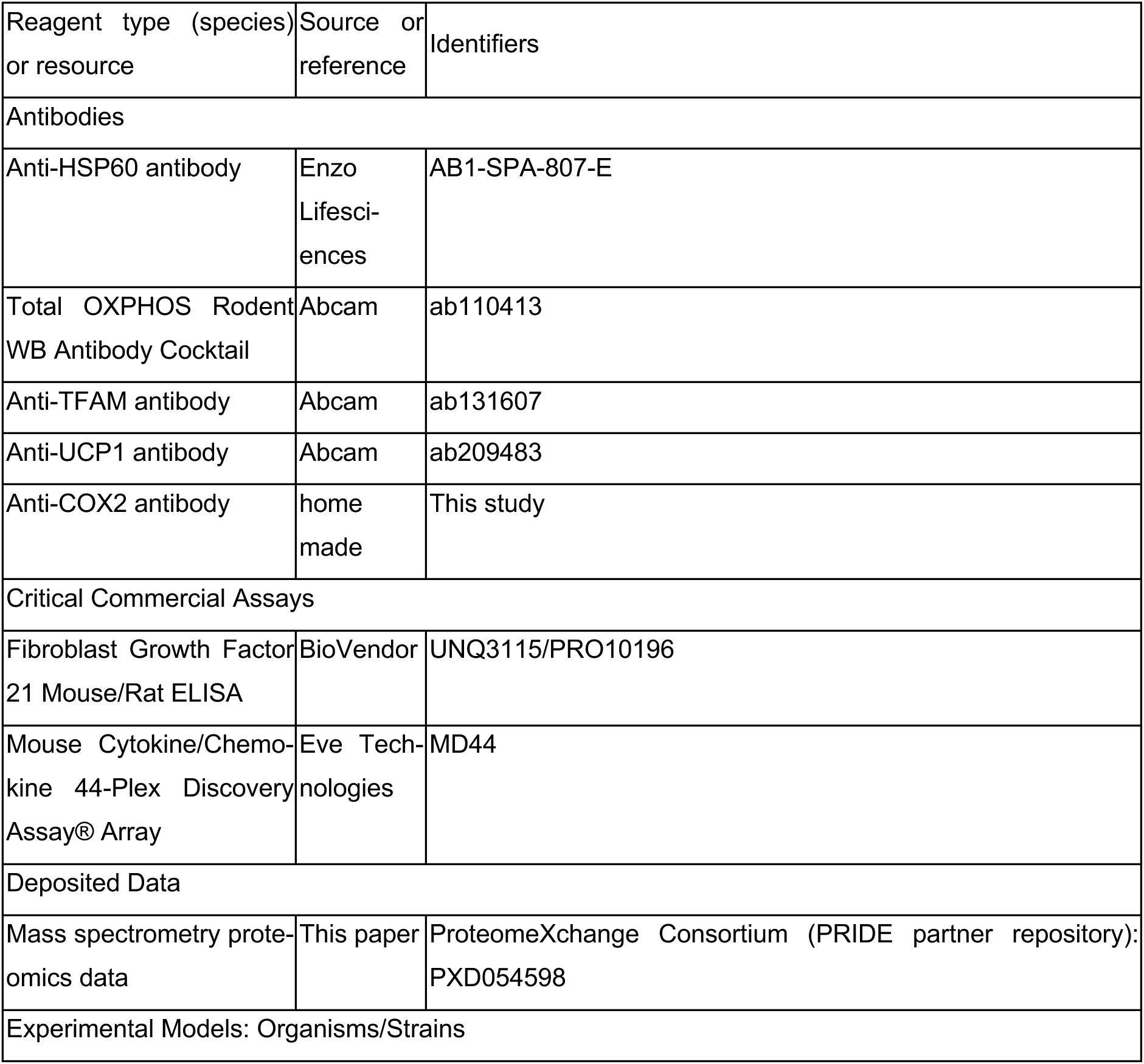
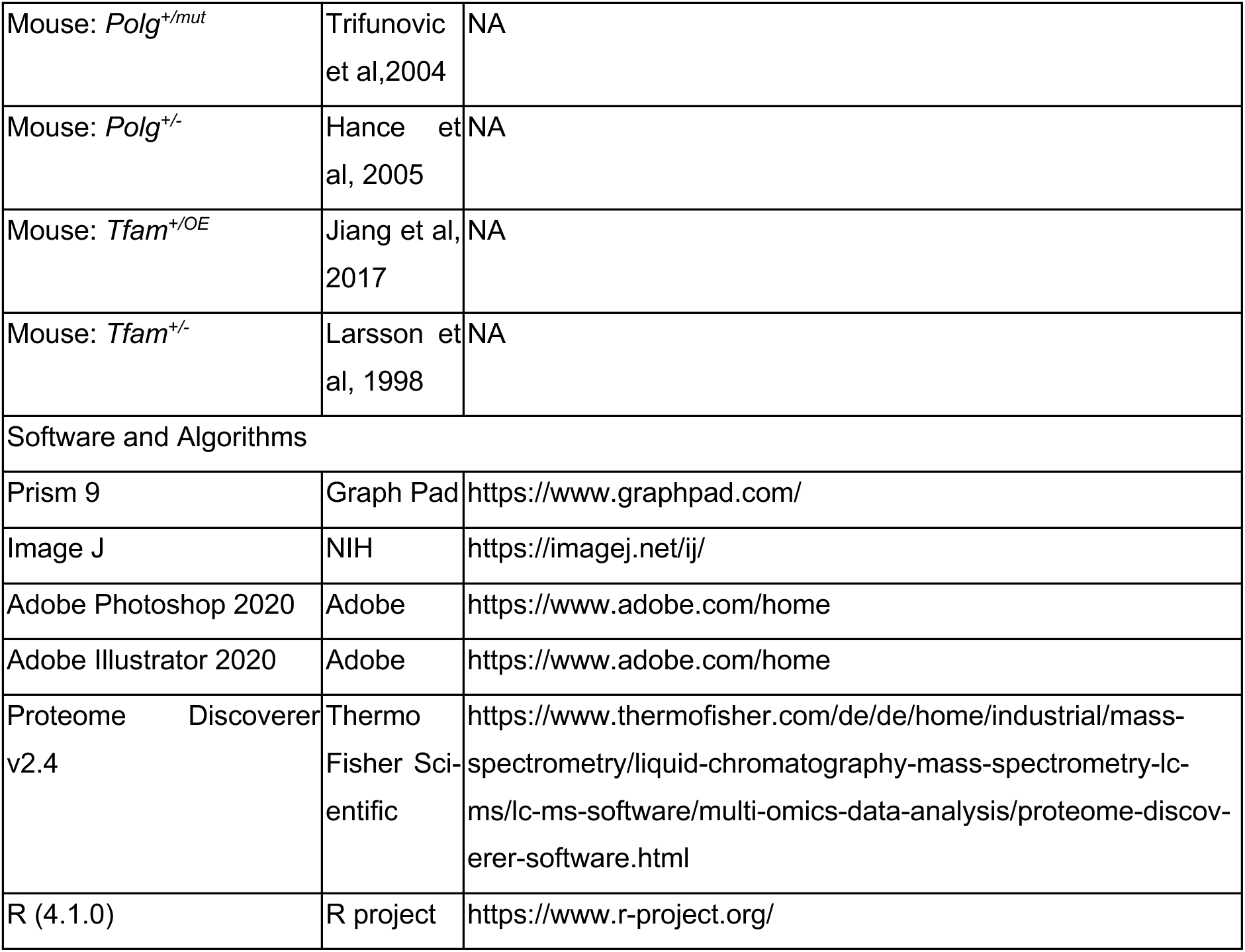

| Reagent type (species) or resource | Source or reference | Identifiers |
| --- | --- | --- |
| Antibodies |  |  |
| Anti-HSP60 antibody | Enzo Lifesciences | AB1-SPA-807-E |
| Total OXPHOS Rodent WB Antibody Cocktail | Abcam | ab110413 |
| Anti-TFAM antibody | Abcam | ab131607 |
| Anti-UCP1 antibody | Abcam | ab209483 |
| Anti-COX2 antibody | home made | This study |
| Critical Commercial Assays |  |  |
| Fibroblast Growth Factor 21 Mouse/Rat ELISA | BioVendor | UNQ3115/PRO10196 |
| Mouse Cytokine/Chemokine 44-Plex Discovery Assay® Array | Eve Technologies | MD44 |
| Deposited Data |  |  |
| Mass spectrometry proteomics data | This paper | ProteomeXchange Consortium (PRIDE partner repository): PXD054598 |
| Experimental Models: Organisms/Strains |  |  |
| Mouse: *Polg*<sup>+/*mut*</sup> | Trifunovic et al, 2004 | NA |
| Mouse: *Polg*<sup>+/-</sup> | Hance et al, 2005 | NA |
| Mouse: *Tfam*<sup>+/*OE*</sup> | Jiang et al, 2017 | NA |
| Mouse: *Tfam*<sup>+/-</sup> | Larsson et al, 1998 | NA |
| Software and Algorithms |  |  |
| Prism 9 | Graph Pad | https://www.graphpad.com/ |
| Image J | NIH | https://imagej.net/ij/ |
| Adobe Photoshop 2020 | Adobe | https://www.adobe.com/home |
| Adobe Illustrator 2020 | Adobe | https://www.adobe.com/home |
| Proteome Discoverer v2.4 | Thermo Fisher Scientific | https://www.thermofisher.com/de/de/home/industrial/mass-spectrometry/liquid-chromatography-mass-spectrometry-lc-ms/lc-ms-software/multi-omics-data-analysis/proteome-discoverer-software.html |
| R (4.1.0) | R project | https://www.r-project.org/ |

### Mouse work

The heterozygous *Polg^+/mut^* mutator mice, the heterozygous *Polg^+/-^*, the *TFAM BAC* (*Tfam^+/OE^*) and *Tfam^+/-^* mice were generated previously as described ^12,24,25,46^. Heterozygous *Polg* knockout (*Polg^+/-^*) males were mated to females that carry either the *Tfam^+/OE^* or *Tfam^+/-^* allele, the resulting *Polg^+/-^; Tfam^+/OE^* or *Polg^+/-^; Tfam^+/-^* females were further mated to heterozygous mtDNA mutator (*Polg^+/mut^*) males to generate the four genotypes used in this study (illustrated in Supp Fig 1). Transgenic mice on a pure C57BL/6N background were housed in a 12-hours light/dark cycle in standard individually ventilated cages and fed *ad libitum* with a normal chow diet. Experimental groups included only 35-week-old male animals. The study was approved by the animal welfare ethics committee and performed in accordance with Swedish and European law.

### Tissue isolation

Animals were euthanized by CO_2_ followed by cervical dislocation. Blood was taken by heart puncture, collected in EDTA tubes and centrifuged at 2000 *g* and 4°C for 10 min to separate the plasma. Heart, spleen, liver, testis, colon, and BAT were collected immediately, washed with phosphate-buffered saline (PBS), and a piece of each tissue snap-frozen in liquid nitrogen, and stored at -80°C. For hematoxylin and eosin (H&E) staining, tissues were embedded in O.C.T. compound (Tissue-Tek), frozen in isopentane precooled in liquid nitrogen, and stored at -80°C. For the liver and colon, an additional tissue piece was kept in PBS for subsequent isolation of crude mitochondria.

### Mitochondrial isolation

Liver and colon were homogenized in mitochondrial isolation buffer containing 320 mM sucrose, 1 mM EDTA, and 10 mM Tris-HCl, pH 7.4, supplemented with 0.2% bovine serum albumin (Sigma-Aldrich), EDTA-free complete protease inhibitor cocktail and PhosSTOP tablets (Roche) by using a Teflon pestle (Schuett Biotec). After centrifugation at 1,000x g (swing-out rotor) for 10 min at 4°C, the supernatants were subsequently spun at 10,000xg for 10 min at 4 °C to isolate the mitochondria. Crude mitochondrial pellets were resuspended in a suitable amount of mitochondrial isolation buffer.

### DNA isolation and mtDNA quantification by qPCR

Genomic DNA from snap-frozen heart, spleen, liver, colon, and BAT was isolated using the DNeasy Blood and Tissue Kit (Qiagen), following the manufacturer’s instructions. Quantification of mtDNA copy number was performed in triplicates using 5 ng of DNA using TaqMan Universal Master Mix II and TaqMan probes (Life Technologies). The mtDNA levels were assessed using probes against the mitochondrial genes encoding *Nd1*, *Atp6*, and *Cytb* and normalized to the nuclear gene encoding *18S* rRNA.

### DNA isolation and mtDNA quantification by Southern blot analysis

Genomic DNA from snap-frozen liver was isolated using the Puregene Cell and Tissue Kit (Qiagen), following the manufacturer’s instructions. Southern blot analysis was performed as described previously using CytB to detect mtDNA and 18S rDNA as nuclear loading control ^47^.

For mouse samples, 2ug total genomic DNA was digested with SacI-HF at 37°C overnight and preheated at 93°C for 3 min, followed by cooling on ice before loading onto the gel. After electrophoresis in 0.8% agarose, DNA was depurinated by incubation in 0.25 M HCl for 10 min and incubated in denaturation buffer (0.5 M NaOH and 1.5 M NaCl) twice for 30 min and neutralization buffer [0.5 M tris-HCl (pH 7.4) and 1.5 M NaCl] twice for 30 min. DNA was blotted onto a Hybond N+ nitrocellulose membrane for 72 hours and then cross-linked by exposure to 254-nm ultraviolet, 200 mJ/cm^2^. Next, membranes were hybridized with α-[^32^P]-dCTP-labelled DNA probes to detect mtDNA (CytB) or nuclear 18S rDNA as a loading control. Radioactive signals were visualized using PhosphorImager screens and a Typhoon 7000 FLA (GE Healthcare). Band intensities were quantified using ImageJ software.

### RNA isolation and quantitative reverse transcription PCR

Total RNA from snap-frozen heart, spleen, liver, colon, and BAT was isolated using TRIzol/chloroform extraction method and quantified with a Qubit fluorometer (Life Technologies). After deoxyribonuclease treatment, reverse transcription was performed using the High-Capacity cDNA Reverse Transcription Kit (Applied Biosystems, Life Technologies). RT-qPCR was performed using the TaqMan Universal Master Mix II with TaqMan probes for *Nd1, Atp6, Cytb, Anf, Atf4, Atf5, Mthfd2, Ucp1, Il-5, Ccl2,* and b*-actin*) (Life Technologies). b-*actin* was used as the loading control.

### Western blot

Tissues were homogenized in RIPA buffer (50 mM Tris pH 7.4, 150 mM NaCl, 1% Nonidet P-40, 0.5% DOC, 0.1% SDS) on ice. The supernatant was collected after centrifugation at 10,000*g* for 10min at 4°C. Protein concentration was determined by the BCA assay. After mixing with NuPAGE™ LDS Sample Buffer (Invitrogen), 20 µg of total tissue lysate was loaded into 12% precast gels (Invitrogen) and separated by SDS-PAGE. Protein was transferred onto polyvinylidene difluoride membranes using the iBlot 2 Gel Transfer system (Invitrogen). Immunodetection was performed according to standard procedure using enhanced chemiluminescence (Clarity ECL Western Blotting Substrates, Bio-Rad) and imaged using the Bio-Rad ChemDoc system. Images were exported using the Bio-Rad ImageLab software and quantified using Image J. The following antibodies were used: HSP60 (Enzo Lifesciences AB1-SPA-807-E), Total OXPHOS Rodent WB Antibody Cocktail (ab110413, abcam), TFAM (ab131607, abcam), UCP1 (ab209483, abcam), COX2 (rabbit polyclonal antisera against COX2 generated using recombinant mouse protein).

### TFAM-to-mtDNA ratios calculation

Average TFAM levels were quantified using Image J and normalized to HSP60 levels (n=2). To compare between the different groups, the averaged data was normalized to the wildtype (*Polg^+/+^Tfam^+/+^*). For the mtDNA level, qPCR data from the ND1 probe was used (n>=5). For group comparisons, the averaged data was normalized to the wildtype (*Polg^+/+^Tfam^+/+^*). Eventually, TFAM-to-mtDNA ratios were calculated by dividing the wildtype normalized TFAM levels by the wildtype normalized mtDNA levels.

### Histochemistry

For H&E staining, the spleen was cryosectioned at −20°C (10 µm section; Cryostar NX70-Thermo Fisher) onto Polysine coated slides (VWR 631-0107) and stored at −80°C until use. H&E staining was performed according to standard procedure. In short, slides were brought to room temperature, washed once in water and stained with hematoxylin solution (ab220365, abcam) for 10 min. The slides were washed in water twice and once in 0.1% sodium bicarbonate. After washing in 96% ethanol, the slides were stained in Eosin Y solution (0.25%) for 5 s. The slides were washed in ethanol, dehydrated and mounted for bright-field microscopy.

### OXPHOS activity measurements

Spectrophotometrically assessment of OXPHOS enzyme activities was performed as previously described^48^. In brief, 500 µg isolated mitochondria were resuspended in 100 µl resuspension buffer (250 mM sucrose, 15 mM KH_2_PO_4_, 2 mM MgAc_2_, 0.5 mM EDTA, 0.5 g/l HSA pH 7.2) and stored as 10 µl aliquots at −80 °C. All assays were performed using an Indiko automated photometer (Thermo Fisher Scientific) fitted with filters for 340, 405, 550, and 600 nm (bandwidth ±5 nm) at 37 °C.

For activity measurements of NADH:coenzyme Q reductase (complex I) and NADH:cytochrome c reductase (complex I + III), samples were pretreated by adding 590 µl of a solution of 5 mM KH_2_PO_4_, 5 mM MgCl_2_, 0.5 g/l HAS (pH 7.2) to the 10 µl frozen mitochondrial suspension. Within 1 min, another 50 µl of the same solution supplemented with 7.15 g/l saponin was added.

For measurement of complex I activity, 72 µl pretreated mitochondria were incubated for 7 min in a reaction mixture with the a final composition of 50 mM KH_2_PO_4_, 5 mM MgCl_2_, 5 g/l HSA, 0.2 mM KCN, 1.2 mg/l antimycin A, and 0.12 mM coenzyme Q_1_ (pH 7.5). Subsequently, NADH was added to a final concentration of 0.15 mM, and the decrease in absorbance was monitored for 1 min before and after the addition of 2 mg/l rotenone at 340 nm in a final volume of 150 µL. For the rotenone-sensitive activity calculation, an extinction coefficient of 6.81 l/mmol/cm was used. For measurement of complex I + III activity, 6 µL pretreated mitochondria were incubated for 7 min in a reaction mixture with the final composition of 50 mM KH_2_PO_4_, 5 mM MgCl_2_, 5 g/l HAS, 0.2 mM KCN, and 0.12 mM cytochrome c (oxidized form) (pH 7.5). Subsequently, NADH was added to a final concentration of 0.15 mM, and the increase in absorbance was monitored for 1 min before and after the addition of rotenone, 2 mg/L at 550 nm in a final volume of 125 µl.

For activity measurements of succinate dehydrogenase (complex II) and succinate:cytochrome c reductase (complex II + III), samples were pretreated by incubating 10 µl of the frozen mitochondrial suspension for 30 min in 100 µl of 50 mM KH_2_PO_4_, 30 mM succinate, 7.5 mM MgCl_2_, and 0.45 g/l saponin (pH 7.2) at 37 °C. Complex II activity was determined as previously described^49^. 10 µl pretreated mitochondria were incubated for 15 min in a reaction mixture of 20 mM KH_2_PO_4_, 5 mM MgCl_2_, 25 mM succinate, 0.2 mM KCN, 0.05 mM 2,6-dichloroindophenol (DCIP), and 2 mg/l antimycin A (pH 7.5). Subsequently, the blank rate was measured for 1 min followed by Coenzyme Q_1_ addition to a final concentration of 0.05 mM and monitoring the decrease in absorbance at 600 nm for 1 min in a final volume of 150 µL. For activity calculation an extinction coefficient of 22 l/mmol/cm was used.

Measurement of complex II + III activity was performed by measuring the blank rate in 50 mM KH_2_PO_4_, 5 mM MgCl_2_, 5 g/l HSA, 0.2 mM KCN, 30 mM succinate, 2 mg/l rotenone, and 0.12 mM cytochrome c (oxidized form) (pH 7.5). Subsequently, 5 µl pretreated mitochondria were added and the reduction of cytochrome c was monitored for 2 min at 550 nm in a final volume of 150 µl.

For cytochrome c oxidase (complex IV) activity measurement, pretreatment was performed by diluting mitochondria to a concentration corresponding to 100 U/l of citrate synthase in 1 g/l digitonin, and 50 mM KH_2_PO_4_ (pH 7.5). The blank rate was recorded in a solution of 50 mM KH_2_PO_4_, 2 mg/l rotenone, and 0.03 mM cytochrome c (reduced form) (pH 7.5). Subsequently, 10 µl pretreated mitochondria were added and oxidation of cytochrome c was followed for 1 min at 550 nm in a final volume of 250 µl.

Citrate synthase activity was used as mitochondrial marker and determined as previously described^50^. 10 µl frozen mitochondria were pretreated by adding 240 µl of 50 mM KH_2_PO_4_, 1 mM EDTA, 0.1% Triton X-100 (pH 7.5). 20 µl pretreated mitochondria were then incubated in 50 mM Tris, 0.20 mM 5,5′-Dithiobis(2 nitrobenzoic acid) (DTNB), 0.1 mM Acetyl-CoA (pH 8.1) for 5 min. Subsequently, oxaloacetic acid was added to a final concentration of 0.5 mM and the increase in absorbance was monitored at 405 nm for 1 min in a final volume of 250 µL. For activity calculation an extinction coefficient of 13.6 l/mmol/cm was used.

For BN-PAGE, 100 µg of isolated mitochondria were lysed in 50 µl NativePAGE™ Sample Buffer (Invitrogen) containing 1% (w/v) digitonin (Calbiochem). Samples were incubated for 10 min at 4°C and centrifuged at 17,000 g for 30 min. After centrifugation, supernatants were collected and protein concentration was measured using BCA assay. 1/10 volume of 5% Coomassie blue G-250 (Invitrogen BN2004) was added to the supernatant. 15 µg protein was loaded on 3-12% Bis-Tris NativePAGE gels (Invitrogen) and run according to the manufacturer’s instructions.

For CI in gel activity measurements, the BN-PAGE gel was incubated in 2 mM Tris/HCl pH 7.4, 0.1 mg/ml NADH (Roche), and 2.5 mg/ml nitrotetrazolium blue for blue staining (Sigma) for about 10 minutes. CIV in gel activity was determined by incubating the BN-PAGE gels in 10 ml of 0.05 mM phosphate buffer pH 7.4, 25 mg 3.3’-diamidobenzidine tetrahydrochloride (DAB), 50 mg Cyt C, 3.75 g Sucrose and 1 mg Catalase for approximately 1 h.

### Immune profiling

For cytokine measurements, plasma was sent to Eve Technologies for assessment (Mouse Cytokine/Chemokine 44-Plex Discovery Assay® Array (MD44)).

For FGF21 measurement, the ELISA kit from Biovendor (UNQ3115/PRO10196) was used following the manufacturer’s protocol.

### Quantitative mass spectrometry

Mouse spleen tissue was prepared as described previously^51^ with some modifications. In brief, samples were thawed on ice and 20-25 mg tissue was cut into small pieces and supplemented with 50 µL of 8M urea, 50 µL of 0.2% ProteaseMAX (Promega) in 20% acetonitrile (ACN), 100 mM Tris-HCl, pH 8.5 and 100 mM NaCl and 1 µL pf 100x protease inhibitor (Pierce) before transferring to a prefilled tube containing 400 µm LoBind silica beads. The samples were frozen for a short time before homogenization using a Disruptor Genie at maximal speed on 2800 rpm for 2 min, incubated on ice for 2 min. These steps were repeated five times. The samples were then centrifuged at 13,000*g* for 10 min at 4°C. The supernatant was collected and 100 µL of Tris-HCl was used to wash the beads, which was combined with the supernatant. Proteins were precipitated with 4-fold volumes of chilled acetone before protein concentration was determined by BCA assay (Pierce).mAn aliquot of 30 µg samples was reduced with 2.5 µL of 250 mM dithiothreiotol, alkylated with 3 µL of 500 mM chloroacetic acid, and digested by addition of 0.6 µg of sequencing grade modified trypsin (Promega) and incubation at 37°C for 16 h. The digestion was stopped with 4.5 µL cc. formic acid and incubating the solution at room temperature (RT) for 5 min. The sample was cleaned on a C18 Hypersep plate with 40 µL bed volume (Thermo Fisher Scientific) and dried using a vacuum concentrator (Eppendorf). Biological samples were labeled with TMTpro reagents (Thermo Fisher Scientific) in random order adding 100 µg TMT-reagent in 30 µL anhydrous ACN to each digested sample resolubilized in 85 µL of 50 mM triethylammonium bicarbonate and incubated at RT for 2 h. The labeling reaction was stopped by adding 11 µL of 5% hydroxylamine and incubating at RT for 15 min before combining all 15 biological samples in one vial. The sample was cleaned on a C18 Hypersep plate with 40 µL bed volume (Thermo Fisher Scientific) and dried using a vacuum concentrator (Eppendorf). For fractionation, the TMT-labeled peptides were dissolved in 50 µL of 20 mM ammonium hydroxide and were loaded onto an XBridge bridged ethyl hybrid C18 UPLC column (2.1 mm inner diameter × 250 mm, 2.5 μm particle size, Waters), and profiled with a linear gradient of 5–60% 20 mM ammonium hydroxide in ACN (pH 10.0) over 48 min, at a flow rate of 200 µL/min. The chromatographic performance was monitored by sampling eluate with a UV detector (UltiMate 3000 UPLC, Thermo Fisher Scientific) scanning at 214 nm. Fractions were collected at 30 s intervals into a 96-well plate and combined in 12 samples concatenating 8-8 fractions representing peak peptide elution before drying in vacuum concentrator. Peptides were reconstituted in solvent A (2% ACN, 0.1% FA) and approx. 2 µg samples injected on a 50 cm long EASY-Spray C18 column (Thermo Fisher Scientific) connected to an UltiMate 3000 nanoUPLC system (Thermo Fisher Scientific) using a 90 min long gradient: 4-26% of solvent B (98% ACN, 0.1% FA) in 90 min, 26-95% in 5 min, and 95% of solvent B for 5 min at a flow rate of 300 nL/min. Mass spectra were acquired on a Q Exactive HF hybrid quadrupole Orbitrap mass spectrometer (Thermo Fisher Scientific) ranging from *m/z* 375 to 1500 at a resolution of R=120,000 (at *m/z* 200) targeting 5x10^6^ ions for maximum injection time of 80 ms, followed by data-dependent higher-energy collisional dissociation (HCD) fragmentations of precursor ions with a charge state 2+ to 7+, using 45 s dynamic exclusion. The tandem mass spectra of the top 18 precursor ions were acquired with a resolution of R=60,000, targeting 2x10^5^ ions for maximum injection time of 54 ms, setting quadrupole isolation width to 1.4 Th and normalized collision energy to 33%. Acquired raw data files were analyzed using Proteome Discoverer v2.4 (Thermo Fisher Scientific) with the Mascot Server v2.5.1 (Matrix Science Ltd., UK) search engine against mouse protein database (SwissProt). A maximum of two missed cleavage sites were allowed for full tryptic digestion, while setting the precursor and the fragment ion mass tolerance to 10 ppm and 0.02 Da, respectively. Carbamidomethylation of cysteine was specified as a fixed modification, while TMTpro on lysine and N-termini, oxidation on methionine as well as deamidation of asparagine and glutamine were set as dynamic modifications. Initial search results were filtered with 5% FDR using Percolator node in Proteome Discoverer. Quantification was based on the TMT-reporter ion intensities. The data was analyzed using R (4.1.0) using the preprocessCore^52^ and pheatmap packages.

### Quantification and statistical analysis

The sample size is indicated in the figure legends and included at least five mice where statistical evaluation was performed. ImageJ was used for quantification. Statistical analysis and generation of graphs were performed with GraphPad Prism v9 software except for quantitative mass spectrometry data which was analyzed and plotted using R as described above. Statistical comparisons were performed using one-way analysis of variance (ANOVA), and post hoc analysis was conducted with Dunnett’s multiple comparisons test. Values of *P* < 0.05 were considered statistically significant. Images were processed with Adobe Photoshop 2020 and schematics were created with Adobe Illustrator 2020.

### Data and code availability

The mass spectrometry proteomics data have been deposited to the ProteomeXchange Consortium via the PRIDE^53^ partner repository with the dataset identifier PXD054598. Proteomics data was analyzed using standard packages and did not generate any new code. Any additional information required to reanalyze the data reported in this paper is available from the lead contact upon request.

## Author contribution

L.S.K. and G.G. designed the experiment, performed experimental work and analyzed the data. G.R., R.F., M.M., and R.W. helped with the experiments. A.V. performed proteomics analysis. C.K. helped to plan and supervise the experiments, L.S.K, G.G. and N.G.L. wrote the manuscript. L.S.K. and N.G.L. conceived and supervised the study.

## Data and materials availability

All data needed to evaluate and conclude the findings in this paper are present in the paper.

---

**Supplementary figure 1:**
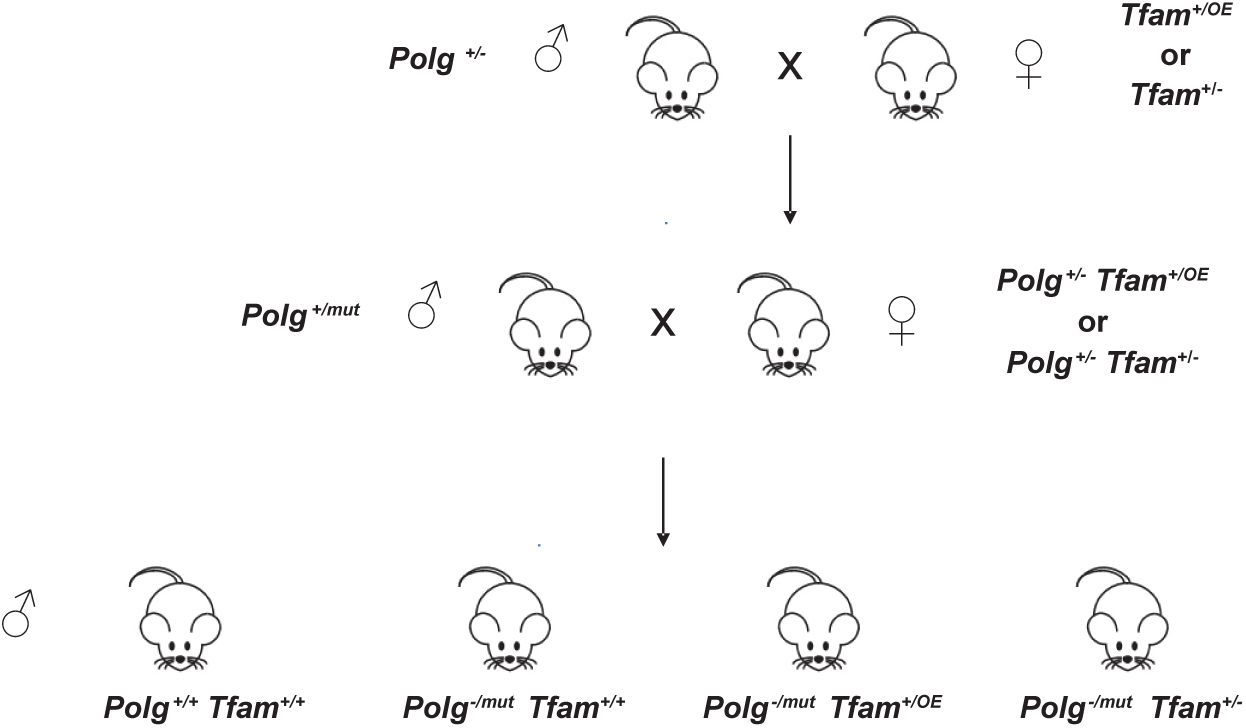
Mating strategy to generate mtDNA mutator mice in combination with the Tfam^+/OE^ or Tfam^+/-^ alleles.

**Supplementary figure 2:**
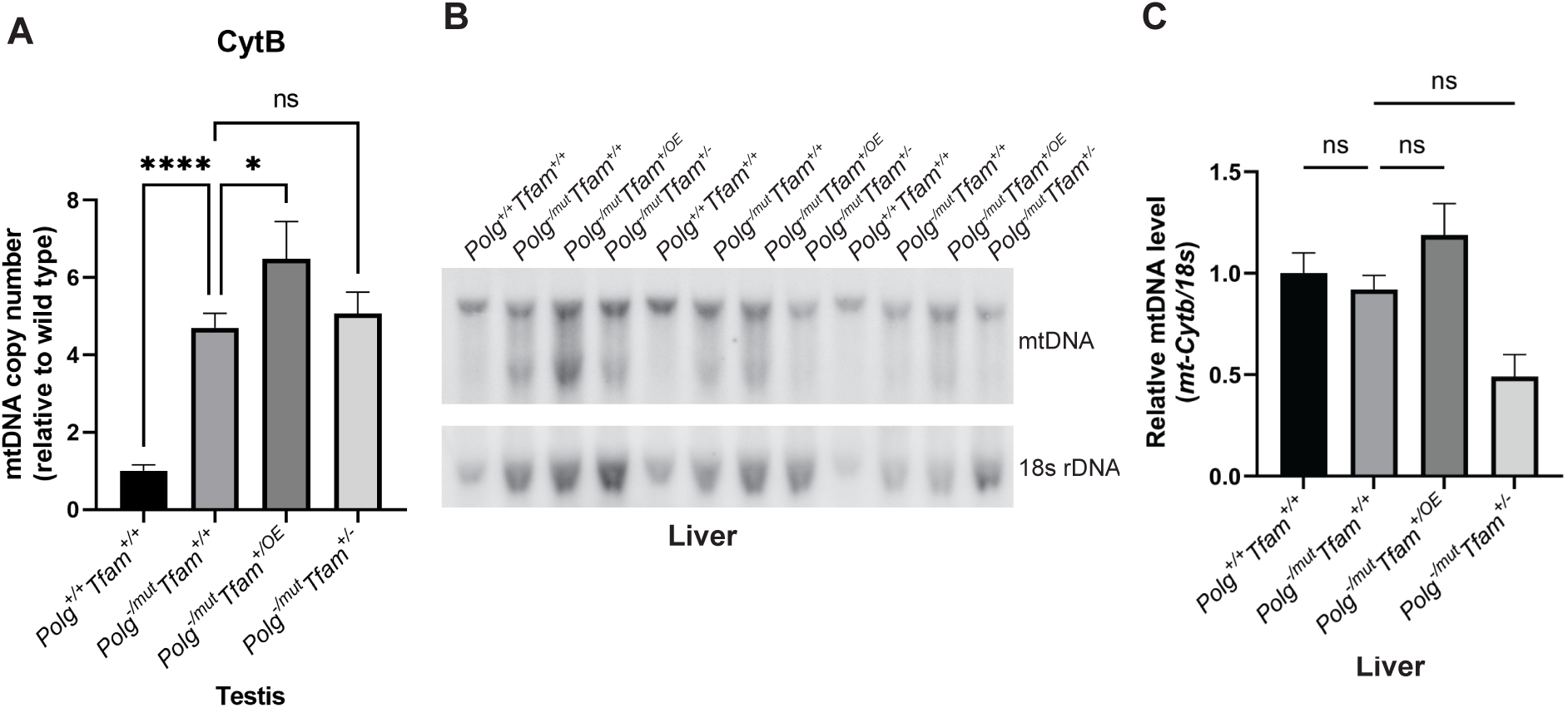
Quantification of mtDNA levels by qPCR in testis and and by Southern blot analysis in liver. A) Relative mtDNA copy number quantification (*Cytb*/*18S*) by qPCR in testis. n≥5. Data are represented as mean ± SEM; *p< 0.05; **p< 0.01; ***p< 0.001; ns: non-significant. B) Southern blot analysis of SacI-digested DNA derived from the liver. mtDNA was quantified by radiolabeling with a specific probe against *Cytb*, nuclear DNA was probed with a specific probe against *18S* rDNA. C) Relative Southern blot quantification (mtDNA/*18S* rDNA); ns: non-significant.

**Supplementary figure 3:**
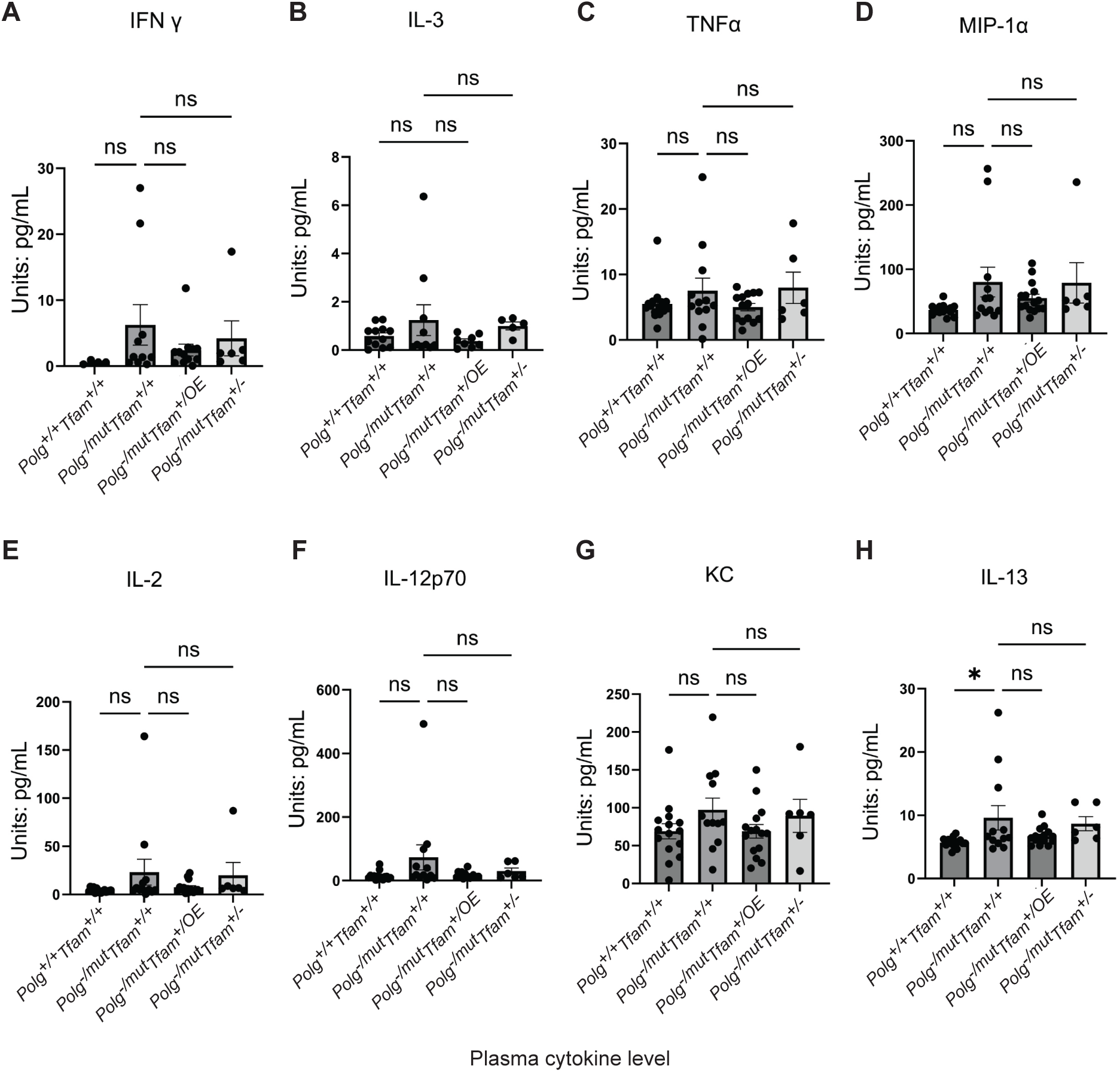
Cytokine quantification in plasma measured by the Mouse Cytokine/Chemokine 44-Plex Discovery Assay. A) Interferon Gamma (IFNγ). B) Interleukin 3 (IL-3). C) Tumor Necrosis Factor Alpha (TNFα). D) Macrophage Inflammatory Protein-1 Alpha (MIP-1α). E) Interleukin 2 (IL-2). F) Interleukin 12p70 (IL-12p70). G) C-X-C Motif Chemokine Ligand 1 (KC). H) Interleukin 13 (IL-13). n ≥6. Data are represented as mean ± SEM; *p< 0.05; **p< 0.01; ***p< 0.001; ns: non-significant.

**Supplementary figure 4:**
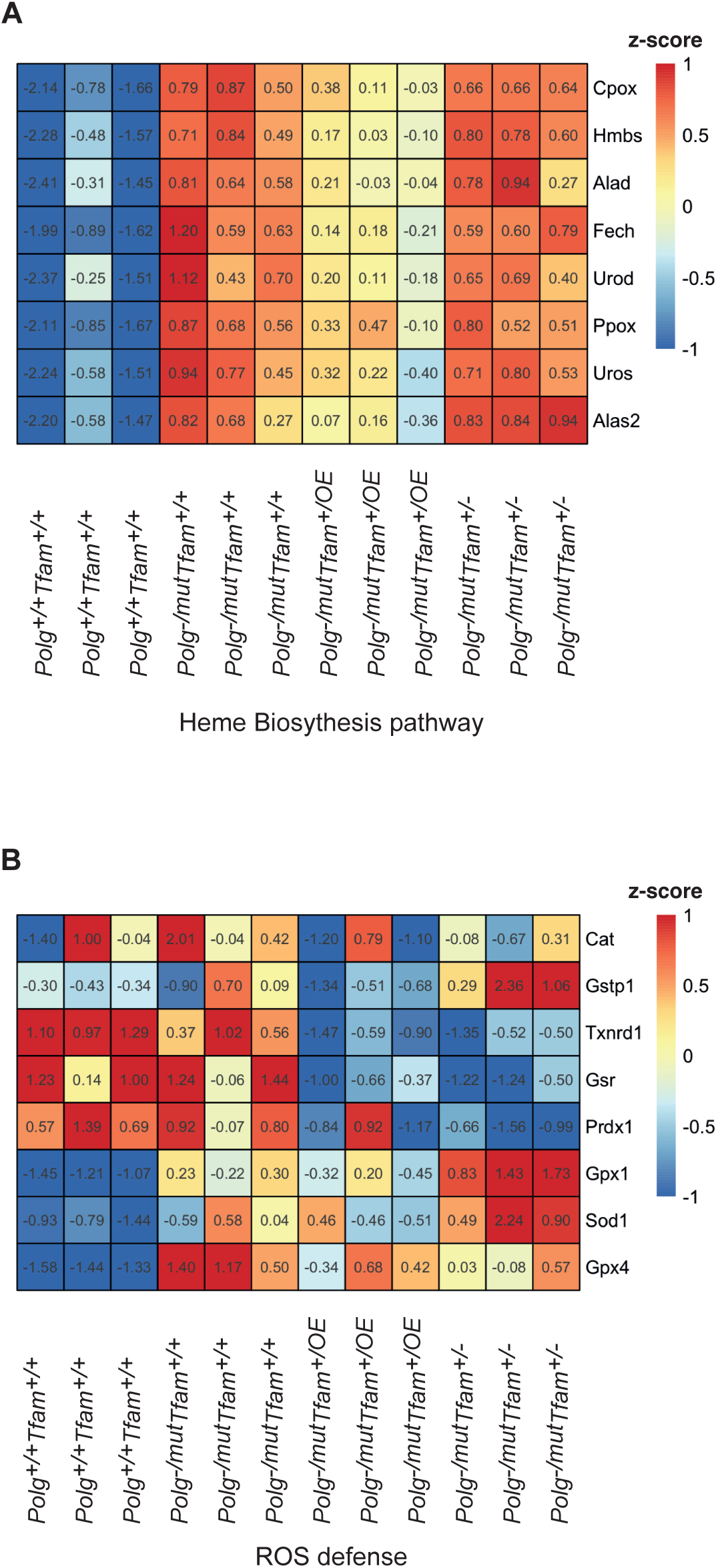
Differential expression analysis of quantitative proteomic data from the spleen. A) Heatmap of proteins involved in the heme biosynthesis pathway. B) Heatmap of proteins involved in ROS defense. Color indicates the z-score.

